# Population ecology and potential biogeochemical impacts of ssDNA and dsDNA soil viruses along a permafrost thaw gradient

**DOI:** 10.1101/2023.06.13.544858

**Authors:** Gareth Trubl, Simon Roux, Mikayla A. Borton, Arvind Varsani, Yueh-Fen Li, Christine Sun, Ho Bin Jang, Ben J. Woodcroft, Gene W. Tyson, Kelly C. Wrighton, Scott R. Saleska, Emiley A. Eloe-Fadrosh, Matthew B. Sullivan, Virginia I. Rich

## Abstract

Climate change is disproportionately warming northern peatlands, which may release large carbon stores via increased microbial activity. While there are many unknowns about such microbial responses, virus roles are especially poorly characterized with studies to date largely restricted to “bycatch” from bulk metagenomes. Here, we used optimized viral particle purification techniques on 20 samples along a highly contextualized peatland permafrost thaw gradient, extracted and sequenced viral particle DNA using two library kits to capture single-stranded (ssDNA) and double-stranded (dsDNA) virus genomes (40 total viromes), and explored their diversity and potential ecosystem impacts. Both kits recovered similar dsDNA virus numbers, but only one also captured thousands of ssDNA viruses. Combining these data, we explored population-level ecology using genomic representation from 9,560 viral operational taxonomic units (vOTUs); nearly a 4-fold expansion from permafrost-associated soils, and 97% of which were novel when compared against large datasets from soils, oceans, and the human gut. *In silico* predictions identified putative hosts for 44% (4,149 dsDNA + 17 ssDNA) of the identified vOTUs spanning 2 eukaryotic, 12 archaeal, and 30 bacterial phyla. The recovered vOTUs encoded 1,684 putative auxiliary metabolic genes (AMGs) and other metabolic genes carried by ∼10% of detected vOTUs, of which 46% were related to carbon processing and 644 were novel. These AMGs grouped into five functional categories and 11 subcategories, and nearly half (47%) of the AMGs were involved in carbon utilization. Of these, 112 vOTUs encoded 123 glycoside hydrolases spanning 15 types involved in the degradation of polysaccharides (e.g., cellulose) to monosaccharides (e.g., galactose), or further monosaccharide degradation, which suggests virus involvement in myriad metabolisms including fermentation and central carbon metabolism. These findings expand the scope of viral roles in microbial carbon processing and suggest viruses may be critical for understanding the fate of soil organic carbon in peatlands.

## Introduction

Soil provides a foundation for a wide range of life on Earth, serves as both a source and a sink of greenhouse gases, and has a key role in the exchange of energy on Earth. Soils and their biodiversity are vulnerable to climate change, especially at northern latitudes where upwards of 50% of global soil carbon (C) is stored in permafrost (McGuire et al. 2018; Schuur et al. 2018), making it the largest terrestrial C stock. Permafrost is thawing at a global rate of ∼1 cm depth/year (Åkerman and Johansson 2008), exposing this large C reservoir to microbial degradation. While the fate of this C remains unknown, a synthesis of previous warming experiments saw considerable C loss from northern latitude soils (Crowther et al. 2016), and the Intergovernmental Panel on Climate Change has highlighted the fate of this C as an important knowledge gap limiting climate change model predictions (Shukla et al. 2019).

Microbes are key drivers in the fate of thawing and thawed permafrost carbon. Microbes degrade complex macromolecules into more labile monomers that either feed C metabolisms, such as fermentation, respiration, and methanogenesis, thereby causing gaseous C loss from the system, or sorb to the soil matrix (Hultman et al. 2015; Woodcroft et al. 2018). As a result, numerous studies seek to understand microbiomes in permafrost ecosystems — largely via culture-independent techniques like paired bulk metagenomes and metatranscriptomes, which has led to a better mechanistic understanding of microbial ecophysiology in these habitats (Hultman et al. 2015; Mackelprang et al. 2016; Ward et al. 2017; Mondav et al. 2017; Woodcroft et al. 2018; Singleton et al. 2018; Wu et al. 2021; Hough et al 2020; Wilson et al. 2022). A synthesis of these data highlights soil moisture content, redox chemistry, sunlight, temperature, and overlying vegetation as key drivers of the microbial response to permafrost thaw. Notably, in at least one case, the abundance of a single microbe could predict ecosystem C fluxes better than any other variable measured (McCalley et al. 2014), which emphasizes the value of illuminating the “microbial black box” for understanding ecosystem functioning. Evidence of the system-level magnitude of microbial impacts is accumulating, and while we currently are unable to predict post-thaw microbial assembly patterns, viruses will undoubtedly play a role.

Viruses are an essential part of soil microbiomes and may be critical for understanding the fate of soil organic carbon. In marine ecosystems, viruses are responsible for lysing approximately one-third of microbes each day, releasing carbon and nutrients on a global scale (Fuhrman JA. 1999; Suttle CA. 2005; Suttle CA. 2007; Breitbart M. 2012). In addition to cell lyses, viruses influence C cycling by metabolically rewiring their hosts, which can result in completely different ‘ecosystem outputs’ for virus infected cells as compared to their uninfected counterparts (Guidi et al. 2016; Gregory et al. 2019; Howard-Varona et al. 2020; Dominguez-Huerta et al. 2022). The metabolic rewiring may include the expression of virus-carried auxiliary metabolic genes (AMGs) that specifically augment host metabolism towards a larger ecosystem impact (Zimmerman et al. 2020). However, translating the roles of marine viruses to soil viruses is difficult, partially due to the extensive microbial richness in soils that obscures the detection of virus genomes (Howe et al. 2014; Thompson et al. 2017). In previous efforts to mine the virus signal in bulk metagenome soil samples, less than 2% of assembled reads targeted viruses (Goordial et al. 2017; Trubl et al. 2018; Emerson et al. 2018). Recent studies have reduced microbial richness by targeting specific microbes with isotopically labeled substrates, allowing increased understanding of virus effects on microbe ecophysiology (Lee et al. 2021; Starr et al. 2021; Trubl et al. 2021; Lee et al. 2022a; Barnett and Buckley 2023; Nicolas et al. bioRxiv), but resolution on virus genomes (i.e., the richness of virus genomes detected and their coverage) in soils remains low.

The advent of virus-targeted metagenomic surveys (better known as viromes) physically separates virus-like particles (VLPs) from the soil matrix with chemical and physical methods, followed by size filtration, VLP concentration and purification, and then DNA extraction (reviewed in Trubl et. al. 2020) These additional steps provide increased resolution on viruses by targeting the estimated 10^7^–10^10^ VLPs per gram of soil (reviewed in Williamson et al. 2017) and removing larger organisms that recruit sequencing data (Trubl et al. 2018). Therefore, viromes capture a higher per-sample vOTU recovery compared to bulk metagenomes (Santos-Medellín, et al. 2021; ter Horst, et al. 2021) allowing for a more comprehensive view of DNA soil viral diversity and in-depth analyses into their evolution and ecology.

Hints of viral roles are starting to emerge, particularly in the natural thawing permafrost gradient of the long-term field site of Stordalen Mire, northern Sweden (Bolduc et al. 2020). At this site, microbial community structure and metabolisms (encoded and expressed) dramatically shift with permafrost thaw (Mondav et al. 2017, Woodcroft et al. 2018), as do viruses recovered from a survey of 178 bulk soil metagenomes (Emerson et al. 2018), 12 size-fractioned metagenomes (Emerson et al. 2018), and 13 viromes (Trubl et al. 2018; Trubl et al. 2019). These viral communities are highly diverse, endemic to their habitat, are predicted to infect key C-cycling microbes and carry glycoside hydrolases, considered AMGs, capable of influencing host C cycling. Notably, six of the viromes captured single-stranded (ss)DNA viruses in addition to double-stranded (ds)DNA viruses and while ssDNA viral ecology was not assessed, the ssDNA viruses recruited a very small proportion of the viromic reads (Trubl et al. 2019). Despite the advances in soil viromics, there are still very few soil viromes from permafrost habitats and a knowledge gap remains in understanding the breadth of C metabolism that viruses modulate and how it changes with permafrost thaw.

Here we applied previously optimized sampling (Trubl et al. 2016, Trubl et al. 2019) to purify virus particles from 20 samples across multiple depths along the Stordalen Mire permafrost thaw gradient over two consecutive years. We compared two existing library preparation techniques, capturing both ssDNA and dsDNA viruses, used optimized bioinformatics methods (Roux et al. 2019a; Pratama et al. 2021) to detect and characterize virus populations, connect the virus populations to putative microbial hosts, and characterize virus-microbe interactions, to provide a baseline understanding of virus diversity and ecosystem impacts. Our optimized sample-to-ecology pipeline had a 10-fold improvement of vOTU recovery (∼30 vOTUs/Gbp of virome versus ∼3 vOTUs/Gbp of virome) compared to our previous method (Trubl et al. 2018).

## Methods and Materials

### Generating viromes

Soil cores from Stordalen Mire were collected in triplicate during the third week of July 2016 and 2017, immediately frozen with liquid nitrogen, shipped to Columbus, Ohio, and stored at −80°C until processed in October 2017 (Supplementary Table 1). Viruses were resuspended from twenty samples from depths 10–14 and 30–34 cm in 2016 samples, and from depths 1–5, 10–14, 20–24, and 30–34 cm in 2017, using a previously optimized method (Fig. 1; Trubl et al. 2016; Trubl et al. 2019). Briefly, 10 ml of a 1% potassium citrate buffer amended with 10% phosphate buffered saline and 150 mM magnesium sulfate was added to 10 ± 0.5 g peat samples in triplicate. The viruses were physically desorbed from soil colloids using vortexing for 1 min and manual shaking for 30 seconds then shaking the tubes at 400 rpm for 15 min at 4°C, and finally, the tubes were centrifuged for 20 min at 4700 ×*g* at 4°C to pellet debris, and the supernatant was transferred to a new tube. The resuspension steps above were repeated two more times and the supernatants were combined and filtered through a 0.2 µm polyethersulfone membrane filter. The viral DNA was extracted using DNeasy PowerSoil extraction kit (Qiagen, Hilden, Germany, cat# 12888), using a heat lysis step: 10 min at 70°C, vortex for 5 seconds and then continue incubating at 70°C for another 5 min. The extracted DNA was quantified using a Qubit-fluorometer (Invitrogen) and the 20 samples were split into duplicates making 40 samples. The Accel-NGS 1S Plus DNA library kit was previously shown to quantitatively capture ssDNA and dsDNA viruses, but to evaluate how well it captures dsDNA viruses, we compared it to a traditionally used Nextera XT DNA library kit. Therefore, half of the samples had libraries prepared using the Accel-NGS kit (Washtenaw County, Michigan) and the other half using the Nextera XT kit (Illumina, San Diego, CA; cat# FC-131-1024). The 40 virome libraries were sequenced with Illumina HiSeq-1TB, run type 2 x 151 (more information in Supplementary Table 2). The Accel NGS 1S plus was the designation at Swift BioSciences which is no longer in business. The comparable kit for the new manufacturer (Integrated DNA technologies) is "xGen ssDNA and low-input". Recently, SRSLY was proposed and could provide another option for quantitative ssDNA/dsDNA viromics.

**Figure 1.**
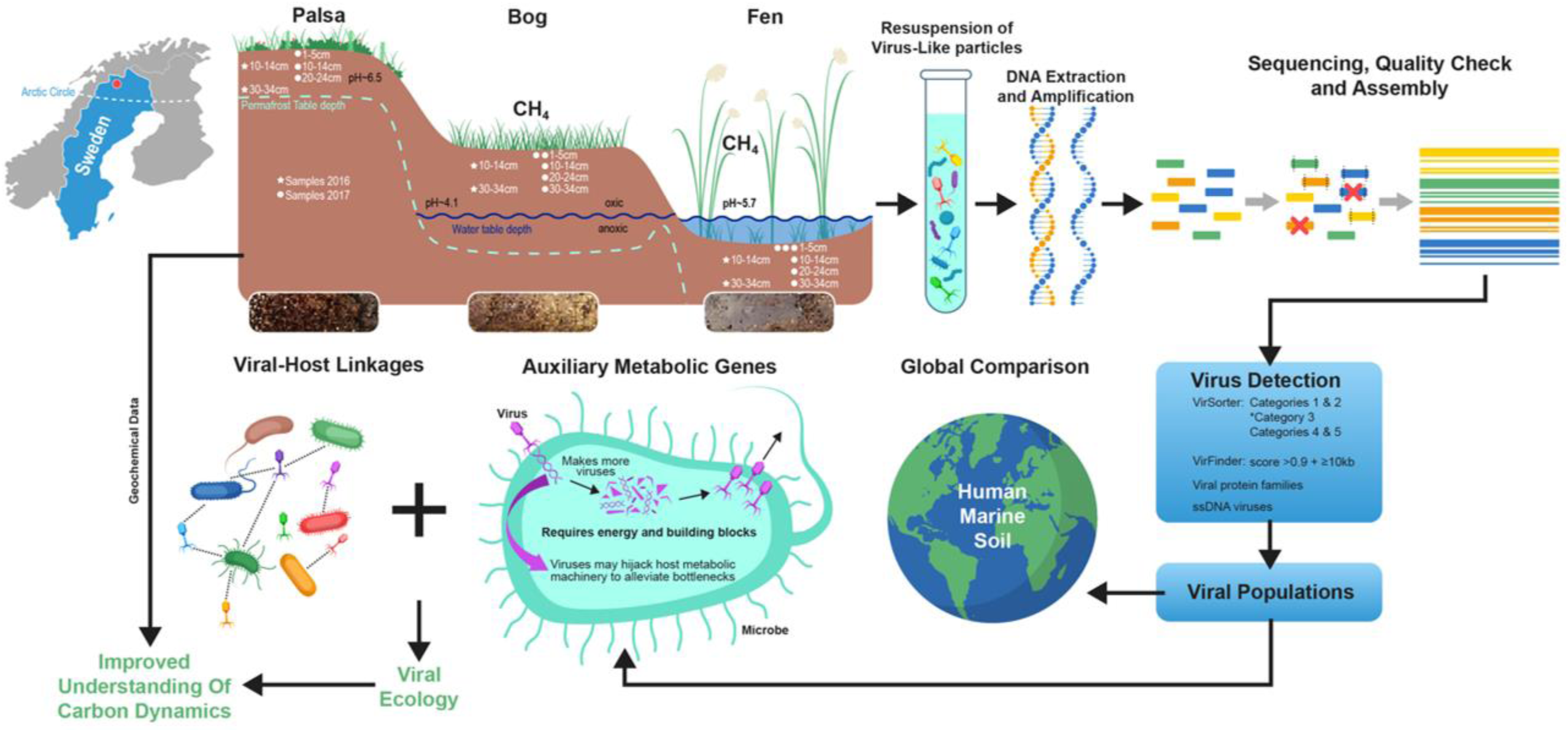
Overview of sampling and methods used to generate viromes. The study site Stordalen Mire (red dot, upper left panel) lies above the Arctic Circle in northern Sweden, and spans three natural permafrost thaw stages (palsa, bog, and fen, with images of representative cores shown below). Soil cores were collected in July 2016 and 2017, with two depths subsampled for viruses in 2016 cores and four subsampled in 2017 cores, for a total of 20 samples for viromes. To achieve the study goals (indicated in green font), best practices in virome recovery, sequencing, and analysis were applied as shown.

### Viral sequence identification and relative abundance estimation

All 40 viromes underwent virus-specific bioinformatic analyses, with the goals of optimizing assembly to aid in virus discovery and increasing coverage from low input libraries (Roux et al. 2019a; Supplementary Table 2). Briefly, the optimized pipeline included “relaxed” read correction with Tadpole (v. 37.76: https://jgi.doe.gov/data-and-tools/bbtools/) to correct sequencing errors by leveraging kmer frequency along each read (parameters “mode=correct ecc=t prefilter=2″), read deduplication with Clumpify (v37.76: https://jgi.doe.gov/data-and-tools/bbtools/), in which identical reads are identified and only one copy retained (parameters: “dedupe subs=0 passes=2″), and assembly using SPAdes (v. 3.11; Nurk et al. 2013; Nurk et al. 2017) with error correction step skipped, single-cell option and k-mers 21, 33, 55, 77, 99, and 127. Assemblies were evaluated using standard metrics computed with stats.sh from the bbtools suite (https://jgi.doe.gov/data-and-tools/bbtools/; see Roux et al. 2019a). To distinguish viral from microbial contigs, the contigs were processed with VirSorter (virome decontamination mode; Roux et al. 2015) and VirFinder (Ren et al. 2017). Contigs were also compared by BLAST (Altschul et al. 1990) to a pool of possible laboratory contaminants (i.e., phages cultivated in the lab where soil samples were processed: Enterobacteria phage PhiX17, Alpha3, M13, *Cellulophaga baltica* phages, and *Pseudoalteromonas* phages) and to a list of ssDNA virus sequences that can commonly contaminate DNA extraction columns (Naccache et al. 2013). The seven contigs matching these genomes at more than 95% average nucleotide identity (ANI) were removed.

To identify dsDNA viral contigs, we used (*i*) viral contigs ≥10kb from VirSorter categories 1 or 2, category 3 with score ≥30 or a score ≥50 to the Global virome HMMs (Páez-Espino et al. 2017), categories 4 and 5 with ≥60% of the genes identified as prophage, and (*ii*) VirFinder ≥10kb with a score ≥0.9. These dsDNA viral contigs were clustered at 95% ANI and 85% alignment fraction (AF) to create a final vOTU dataset (Roux et al. 2019b) and their quality was assessed with CheckV (Nayfach et al. 2020) (Supplementary Table 3). As is common for soil virome work, only a small amount of DNA was extracted from these samples which pushes the library preparation methods and requires relaxation of read-mapping coverage requirements to see ‘biological’ signals (Paez-Espino et al. 2016; Roux et al. 2017; Trubl et al. 2019). Therefore, reads were mapped back to the nonredundant set of contigs at 90% ANI and two thresholds for the length of contig covered to estimate coverage: 75% contig coverage for our strict threshold as it offers robust coverage of well-sampled populations (Roux et al. 2017), and 10% for our relaxed threshold because it allows for high heterogeneity in populations (Paez-Espino et al. 2016; Paez-Espino et al. 2017; Trubl et al. 2019). Coverage was calculated as the number of base pairs mapped to each read normalized by the length of the contig, and by the total number of base pairs sequenced in the metagenome using Bowtie 2 (Langmead and Salzburg 2012; Roux, et al. 2017). The heat map of the vOTU’s relative abundances was constructed in R v4.2.2 pheatmap v1.0.12. Ecological diversity indices of dsDNA vOTUs were done in R Vegan v. 2.6.4 with Shannon’s index and visualized with ggplot v3.4.0, and accumulation curves with specaccum (method = "random", permutations = 100, conditioned =TRUE, gamma = "jack2“). Linear regression analysis of the recovered viral contigs was done in Excel with the data analyses package and outliers were identified by finding the difference between the first and third quartile of the distribution and multiplying it by 2.2 (Hoaglin, Iglewicz, and Tukey 1986; Hoaglin and Iglewicz, 1987).

A two-step approach was used to specifically identify ssDNA contigs as previously described (Trubl et al. 2019). First, putative complete ssDNA virus genomes were identified based on hmmsearch (Eddy, 2009) hits on circular contigs to ssDNA marker genes from the PFAM database: Phage_F (also known as major capsid protein, MCP) and Viral_Rep (also known as replication associated protein, Rep) domains (HMMER v3; cutoffs: score ≥50 and e-value ≤0.001). The protein sequences similar to the MCP or Rep domains were aligned and used to build new HMM profiles, which were then searched against the set of metagenome-predicted proteins (from VirSorter), using the same hmmsearch approach with similar cutoffs as previously and size thresholds of 1–5kb for Rep/cressdnaviruses and 4–8kb for MCP/microviruses. All *de novo* assembled ssDNA virus contigs were manually checked for terminal redundancy to determine circularity.

The open reading frames (ORFs) of the cressdnaviruses (only complete genomes) were determined using ORFfinder (https://www.ncbi.nlm.nih.gov/orffinder/) couple with manual curation of the splice sites in the replication associated protein (Rep) CDS. The cressdnaviruses were manually annotated based on known ORFs of cressdnavirus Rep and CP. ORFs and annotation of the viruses in the family *Microviridae* were determined using RASTtk (Brettin et al. 2015) with manual refinement. Complete circular genomes of cressdnaviruses and microviruses were linearized staring at the start of the Rep or MCP coding sequences, respectively. These genomes were then binned into vOTUs with 95% ANI using CD-HIT (Fu et al. 2012). Representative genomes from each vOTU were used to map raw trimmed reads from each sample library using BBmap to determine genome coverage.

### Relating vOTUs to other Stordalen Mire vOTUs and global vOTUs

To compare recovered vOTUs to those known globally, we clustered them against previous vOTU data from the same site (2,561; Emerson et al. 2018; Trubl et al. 2018; Trubl et al. 2019), other soils (12,032; Van Goethem et al. 2019; Santos-Medellin et al. 2021; ter Horst et al. 2021; Trubl et al. 2021), marine systems (488,130; Gregory et al. 2019), the human gut (33,242; Gregory et al. 2020), and NCBI reference sequences (14,256; RefSeq v204). The field of viromics is rapidly evolving and every few months new data is being published; therefore, we froze our global analyses in 2021. The vOTUs from all the datasets were clustered at 95% ANI and 85% AF to determine if the vOTUs were unique or shared among the datasets. To quantitatively visualize the data, we generated two static UpSet plots in R (Conway, Lex, and Gehlenborg 2017; databases and other information is in Supplementary Table 4).

The ssDNA viruses were further interrogated by comparing them with previously reported Stordalen Mire ssDNA vOTUs (Trubl et al. 2019), Les Pradeaux Mire ssDNA viruses (Quaiser et al. 2015; Quaiser et al. 2016) and with ssDNA viruses in GenBank. Reps from species of classified cressdnaviruses as well as alphasatellites were clustered with CD-HIT at 60% amino acid identity to generate a representative subsample of these. A representative of each of these clusters, together with 2,587 Reps amino acid sequences of unclassified cressdnaviruses genomes available in GenBank (downloaded October 5, 2021) and 574 Rep amino acid sequences of cressdnaviruses identified in this study were aligned using MAFFT (Katoh and Standley 2013). The MCP amino acid sequences of Microviridae (n=3153) genomes available in GenBank were aligned with 102 MCPs from this study and 8 representatives bullavirus MCPs using MAFFT (Katoh and Standley 2013).

The resulting alignments of the Rep and MCPs was trimmed using trimAl (gap threshold of 0.2; Capella-Gutierrez, Silla-Ma 2009) and used to infer a maximum likelihood phylogenetic tree using IQTree2 (Minh et al. 2020) with Q.pfam+F+G4 substitution model and aLRT branch support. The resulting phylogenetic tree was midpoint rooted and visualized in iTOLv6 and represented as cladogram (Letunic and Bork 2021).

### Linking vOTUs to microbial hosts

We used two approaches based on sequence similarity to predict hosts for the vOTUs. The vOTU representative sequences were compared to the 1,529 Stordalen Mire MAGs (Woodcroft et al. 2018) using BLASTn (Altschul et al. 1990) (options -dust no -perc_identity 70) based on shared genomic content (Edwards et al. 2016), and any vOTU with a bit score ≥50, E-value ≤ 10^-3^, and ≥70% identity across ≥2,500 bp were considered for host prediction (Supplementary Table 5). CRISPR spacer matches were also used to link viruses from this study to the Stordalen Mire MAGs (Woodcroft et al. 2018). To identify CRISPRs *in silico* from the MAGs, we used minced (Bland et al. 2007; https://github.com/ctSkennerton/minced) to extract CRISPR spacers. After removal of spacer sequences with at least one ambiguous base (2 spacers), we identified a total of 3,560 spacer sequences. We searched the 3,560 spacer sequences against all the dsDNA and ssDNA viral population representatives using blastn (Altschul et al. 1990), with the BLASTn-short option and default parameters except for “-qcov_hsp_perc 100”. After conservatively filtering for matches with 100% sequence identity, we detected 42 total spacer-virus matches (Supplementary Table 5). We traced the MAG in which each CRISPR spacer was derived to link the microbial host to its virus. Stacked Bar Plots were made in R using ggplot2 to visualize virus-host connections.

We used an additional integrated approach, iPHoP (Integrated Phage HOst Prediction; Roux et al. 2023), to increase the number of vOTUs connected to host taxa. Briefly, iPHoP attempts to link input phage sequences to a host taxon (typically at a genus or family rank) based on a combination of (i) direct sequence comparison to host genomes and CRISPR spacers, (ii) overall nucleotide composition comparison to host genomes, and (iii) comparison to phages with known hosts. Here, we first used iPHoP v1.2.0 with default parameters to predict hosts for all vOTU representatives based on the default iPHoP database (Sept_2021_pub). Next, we built a custom iPHoP database by adding 1,529 Stordalen Mire MAGs (Woodcroft et al. 2018) to the default GTDB-based iPHoP database (using the “add_to_db” function) and ran a second host prediction on the same vOTU representatives (iPHoP v1.2.0). Host predictions were filtered using a minimum score of 75, each vOTU was connected to the host taxon (at the family rank or if ≥90, the genus rank if available) with the highest score across both iPHoP runs (default and custom database). Host predictions by iPHoP were ignored for vOTUs predicted to infect eukaryotes based on their taxonomic classification (e.g., *Cressdnaviricota*) as these sequences are known to yield false-positive prokaryotic host predictions.

### Predicting viral Auxiliary Metabolic Genes and other metabolic genes from viral contigs

Identification and curation of AMGs was performed following a recently established standard operating protocol (Pratama et al. 2021). Viral contigs and their putative AMGs and metabolic genes were annotated with DRAM-v using the uniref option (--use_uniref; Shaffer et al. 2020). Due to the dependence of DRAM-v on VirSorter for determining confidence, the AMGs and metabolic genes discussed here were from viral contigs called by VirSorter. For each gene on a viral contig that DRAM-v annotated as metabolic, a score, from 1 to 5 (1 being most confident), denotes the likelihood that the gene belongs to a viral genome rather than a degraded prophage region or a poorly defined viral genome boundary (Shaffer et al. 2020). To link these viruses more readily to the C cycle in this system, we included 46 genes as putative AMGs (added to DRAM database) that were highlighted to be important in ecosystem C cycling (Woodcroft et al. 2018; Supplementary Table 6). DRAM-v was run on 4,201 viral contigs were detected using VirSorter that were ≥5 kbp in length and these viral genomes contained 1,684 putative AMGs or metabolic genes with a score of 1–3. To minimize the number of false positives, we considered only genes that were flanked by viral or viral-like genes on either side. For category assignment (e.g., C utilization), AMGs and metabolic genes were assigned first by DRAM-v provided categories, and if no category was provided by DRAM-v, the genes were manually assigned to each category based on annotation (Supplementary Table 6).

## Data availability

Custom perl scripts used in this study are available at https://bitbucket.org/srouxjgi/scripts_pcrlibs_assembly_optimization/src/master/. Reads for the different metagenomes are available on https://genome.jgi.doe.gov/portal/ and the SRA database (https://www.ncbi.nlm.nih.gov/sra). Sample accession numbers are listed in Supplementary Table 1 and GenBank accession numbers for ssDNA vOTUs are listed in Supplementary Table 3.

## Results and Discussion

Stordalen Mire is a long-term study site in northern Sweden that has a mosaic of habitats that span a natural permafrost thaw gradient. Stordalen Mire is easily accessible with built-in infrastructure that offers stable navigation of the terrain with minimal environmental disturbance, eddy covariance flux measurements, and has the Abisko field station nearby, which is well equipped to support research (e.g., laboratories, offices, sleeping quarters). The combination of these qualities has attracted researchers globally and allowed unique long-term experiments to be conducted that provide historical data spanning more than a century. For more than a decade, the IsoGenie project, now the EMERGE Biology Integration Institute, has leveraged the historical data to advance a systems-level understanding of permafrost thaw with matched amplicon, meta-omic, process measurements, and biogeochemical characterization of thaw stages (comprehensive review in Bolduc et al. 2020). We expanded these sampling efforts by collecting 20 peatland samples spanning three stages of permafrost thaw (palsa – intact permafrost, bog – partially thawed permafrost, and fen – fully thawed permafrost) across two years and multiple depths for viromic analyses (Fig. 1). To maximize viral particle recovery, we used protocols optimized for their resuspension (Trubl et al. 2016) and DNA extraction (Trubl et al. 2019) specifically for these soils. We then sought to maximize virus sequences captured by applying two quantitative library approaches that had been previously optimized for dsDNA viruses (Duhaime et al. 2012a; Duhaime et al. 2012b; Solonenko et al. 2013a; Solonenko et al. 2013b; Trubl et al. 2019) or both ssDNA and dsDNA viruses (Roux et al. 2016; Trubl et al. 2019). The 40 resulting libraries were then sequenced to create viromes to quantitatively capture ssDNA and dsDNA viruses as a foundation to study their ecology in the Mire.

### Comparing viral signal recovery from different library methods

To access the viral diversity, two quantitatively proven library methods were used: (1) the Nextera XT kit (here forth Nextera), which amplifies dsDNA (Rinke et al. 2016), and (2) the Accel-NGS 1S plus kit (here forth Accel), which amplifies both ssDNA and dsDNA (Roux et al. 2016; Trubl et al. 2019). We previously developed a pipeline using this dataset combining read deduplication and an assembly algorithm that improved assembly of contigs ≥10 kb and increased recovery of vOTUs by 2-fold for low input metagenomes (Roux et al. 2019a). Applied to the 40 viromes generated here, this pipeline yielded 23,982 dsDNA viral contigs ≥10kb with the Nextera library method, 24,517 dsDNA viral contigs ≥10kb and 4,765 ssDNA viral contigs (see methods) with the Accel library method. Overall, 21,347 dsDNA viral contigs were identified in both datasets (Fig. 2A).

**Figure 2.**
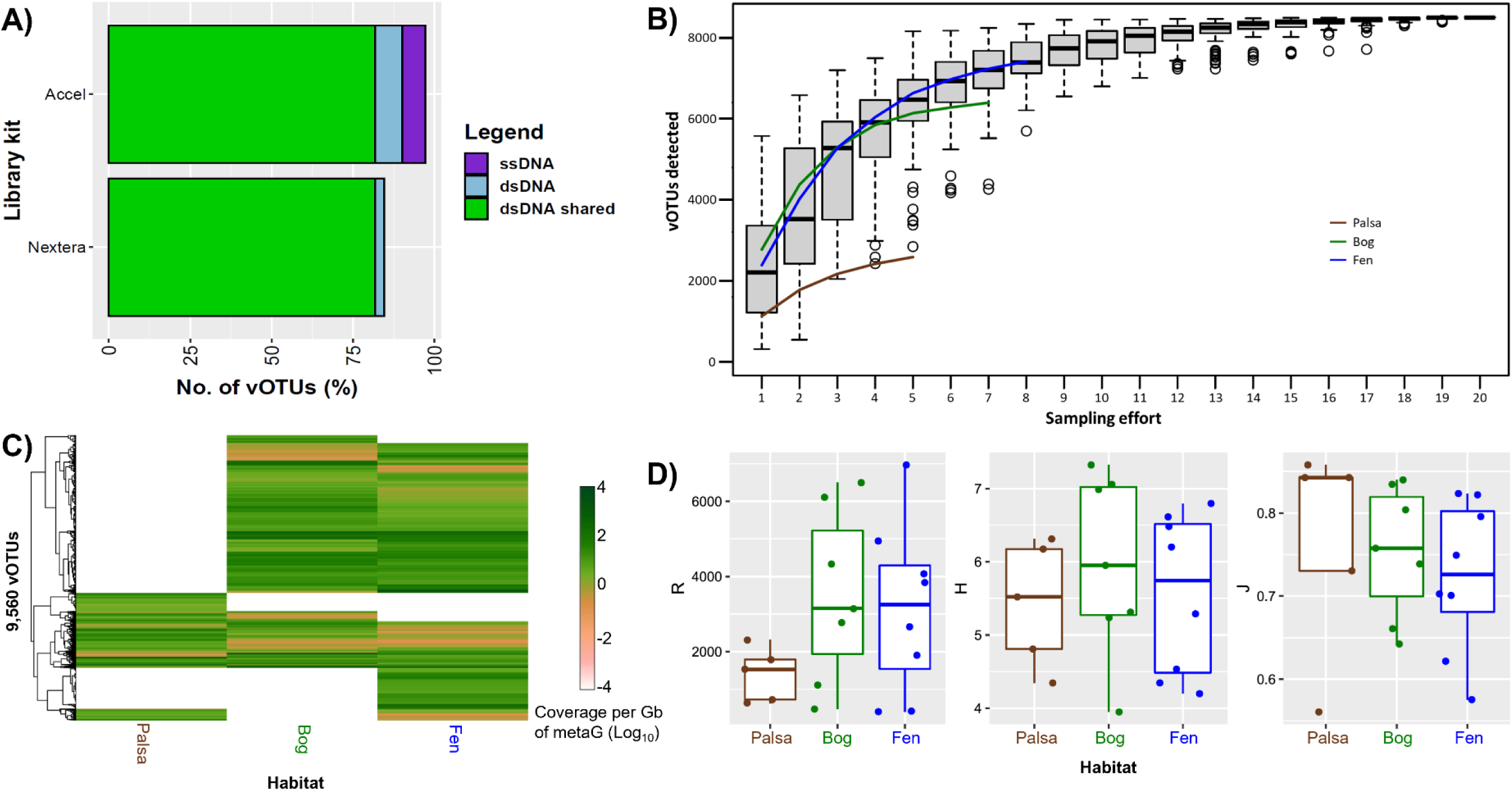
Assessments of viral populations between library kits, and among habitats and samples. (A) A stacked bar chart showing the unique and shared vOTUs for each library method. (B) Accumulation curves of cumulative vOTU richness as sampling effort increased. The gray boxplots represent the average of all the virome data (n=20) and reflect the rate of new vOTUs under continuous sampling (100 permutations). The overlaid lines display the mean cumulative richness per habitat: palsa n=5, bog n=7, and fen n=8. (C) A heatmap showing the coverage of the vOTUs among the three habitats. The vOTUs were clustered using a Bray-Curtis dissimilarity metric based on vOTU normalized abundance. (D) boxplots showing diversity metrics of the vOTUs for each habitat based on vOTU normalized abundance.

We previously noted that the PCR-amplified metagenomes were quantitative when compared to unamplified libraries and reported coverage bias along individual genomes (Roux et al. 2019a). Here, we leveraged the larger dataset of paired soil viromes to further these two library preparation methods in terms of viral contig recovery and potential coverage biases. The number of viral contigs recovered for each sample was mostly consistent between the two methods except for 3 outlier samples (B_Mid_2016, B_Sur_2017, and F_XD_2016; Supplementary Fig. 1A). There were no significant differences in the three outliers’ features (e.g., DNA input, rounds of PCR; ANOVA p >0.05) between the two library datasets, so we proceeded to compare the shared contigs. Comparing the coverage of contigs detected in both kits for each metagenome, revealed the Accel kit performed significantly better at the relaxed threshold (Supplementary Fig. 1B; ANOVA p >0.01) and the kits performed similarly at the strict threshold (Supplementary Fig. 1C). We next compared the coverage per contig detected in both kits and saw larger variation between the kits than variation within each kit for both thresholds (Supplementary Fig. 1D/E), and the Accel kit contigs recruited 74% more basepairs (at both the strict and relaxed thresholds), despite Accel samples having an average of 11% less sequencing effort. Contig coverage biases, specifically unmapped reads, are a common problem with tagmentation (Solonenko et al. 2013b; Jones et al. 2015; Sato et al. 2019; Gunasekera et al. 2021). This problem is often amplified in soils due to the presence of polysaccharides, which are known inhibitors of DNA quantification and enzymatic reactions such as tagmentation which can cause under or over-tagmentation. Illumina recently launched a new version of Nextera called “DNA Prep kit” to address issues associated DNA input by using bead-basedtechnology to accommodate a wide range of DNA amounts, removing the need for DNA quantitation, and reducing under and over-tagmentation (Bruinsma et al. 2018). Future studies are needed to assess varying quantities and compositions of polysaccharides co-eluting with DNA to determine specific associated biases that may still exist.

### Characterization and ecology of viral populations

To explore the ecology of these viruses at the population level, we combined the libraries to create a non-redundant list of viral populations (vOTUs) as reference genomes for all further analyses. This resulted in 8,884 dsDNA vOTUs (3% unique to Nextera and 9% unique to Accel) and 676 ssDNA vOTUs (unique to Accel due to the library preparation selection step) resulting in 9,560 total vOTUs spanning the permafrost thaw gradient (Fig. 2A). We next assessed the quality of the dsDNA vOTUs with CheckV and identified 20 complete genomes, plus 716 high quality, 1,110 medium quality, 5,985 low quality, and 1,053 quality-not-determined viral genomes (Supplementary Table 3). Because CheckV performs best where closely related reference genomes are available and soil virus reference genomes are not widely available, we interpret the large number of ‘not-determined’ and ‘low quality’ dsDNA vOTU genomes to represent an unknown mix of challenges for CheckV and highly fragmented genomes from low coverage soil viromes. Accumulation curves of these dsDNA vOTUs revealed that as the sample size increased, the number of vOTUs initially increased, and then, the curve began to plateau, indicating that the number of vOTUs had reached saturation and that we had enough samples to look for ecological patterns (Fig. 2B). The ssDNA viral genomes were manually inspected for terminal redundancy to determine circularity (see methods) and only complete genomes were retained (913 genomes from 676 vOTUs). These reference genomes represent the first large-scale soil virome dataset of quantitatively amplified ssDNA and dsDNA viruses.

With these reference genomes in-hand, we next evaluated their ecological patterns across this site. Comparing the coverage of vOTUs along the permafrost thaw gradient revealed the fen habitat had the most unique vOTUs (1270; 14%), followed by the palsa habitat (565; 6%) and then bog habitat (246; 3%; Fig. 2C). This result contrasts our previous viromics analysis of these habitats in which bog had the most unique vOTUs, followed by fen and then palsa (Trubl et al. 2018).

Presumably this contradiction is due to the more comprehensive sampling of our current study, which mirrors what is seen in the microbial data with the fen exhibiting unique lineages compared to the pasla and bog (Woodcroft et al. 2018). Previous work from Stordalen Mire also showed that viral community composition was significantly correlated with peat depth (Emerson et al 2018), with the greatest viral abundance in the fen habitat (paralleling microbial abundances), but the highest richness in the bog (unlike the microbes, which are richest in the fen), and a clear pattern of habitat-specific viruses (Emerson et al. 2018; Trubl et al. 2018). In our dataset we saw similar trends with viral richness highest in the bog (Fig. 2D; not significant one-way ANOVA, and Tukey’s test p >0.05), but coverage and richness were higher in the surface and middle depths for each habitat, with the exception of the Palsa surface samples (two-way ANOVA and Tukey’s test p >0.05; Supplementary Fig. 2A), and habitat type was a larger driving factor of viral community composition than depth or year (Supplementary Fig. 2B). We hypothesized that the increased viral richness in the upper depths was likely due to bacterial and archaeal richness also being highest at these depths (Woodcroft et al. 2018).

We further investigated the taxonomy and ecology of the ssDNA vOTUs detected across Stordalen Mire. The 676 ssDNA vOTUs were detected using marker genes – the major capsid protein, or MCP, for viruses in the family *Microviridae* and the replication protein, or Rep, for viruses in the phylum *Cressdnaviricota* (Krupovic et al. 2020) – and identified 572 vOTUs for cressdnaviruses and 102 for microvirus. A comparison of relative abundances between viruses in the phylum *Cressdnaviricota* and family *Microviridae* mirrors their richness, with cressdnaviruses being just over 5-fold more abundant than microviruses (Fig. 3A). A taxonomic breakdown of viruses in the phylum *Cressdnaviricota* revealed that virtually all the ssDNA viruses (98%) were unclassified cressdnaviruses, with just a few viruses in the family *Circoviridae* (n=4; genus *Cyclovirus*), *Genomoviridae* (n=1; genus *Gemycircularvirus*), *Naryaviridae* (n=1; represents a member of a new family) (Fig. 3B). These analyses underestimate the diversity of ssDNA viruses in these soils because it is based solely on the two lineages of viruses captured with marker genes. Nevertheless, detection of ssDNA viruses in environmental samples is difficult due to detection methods using the dsDNA virus features to detect all viruses (Guo et al. 2021) or using of rolling-circle amplification which precludes ecological interpretation (Roux et al. 2016); therefore, marker genes provide the best available strategy to investigate ssDNA viral ecology.

**Figure 3.**
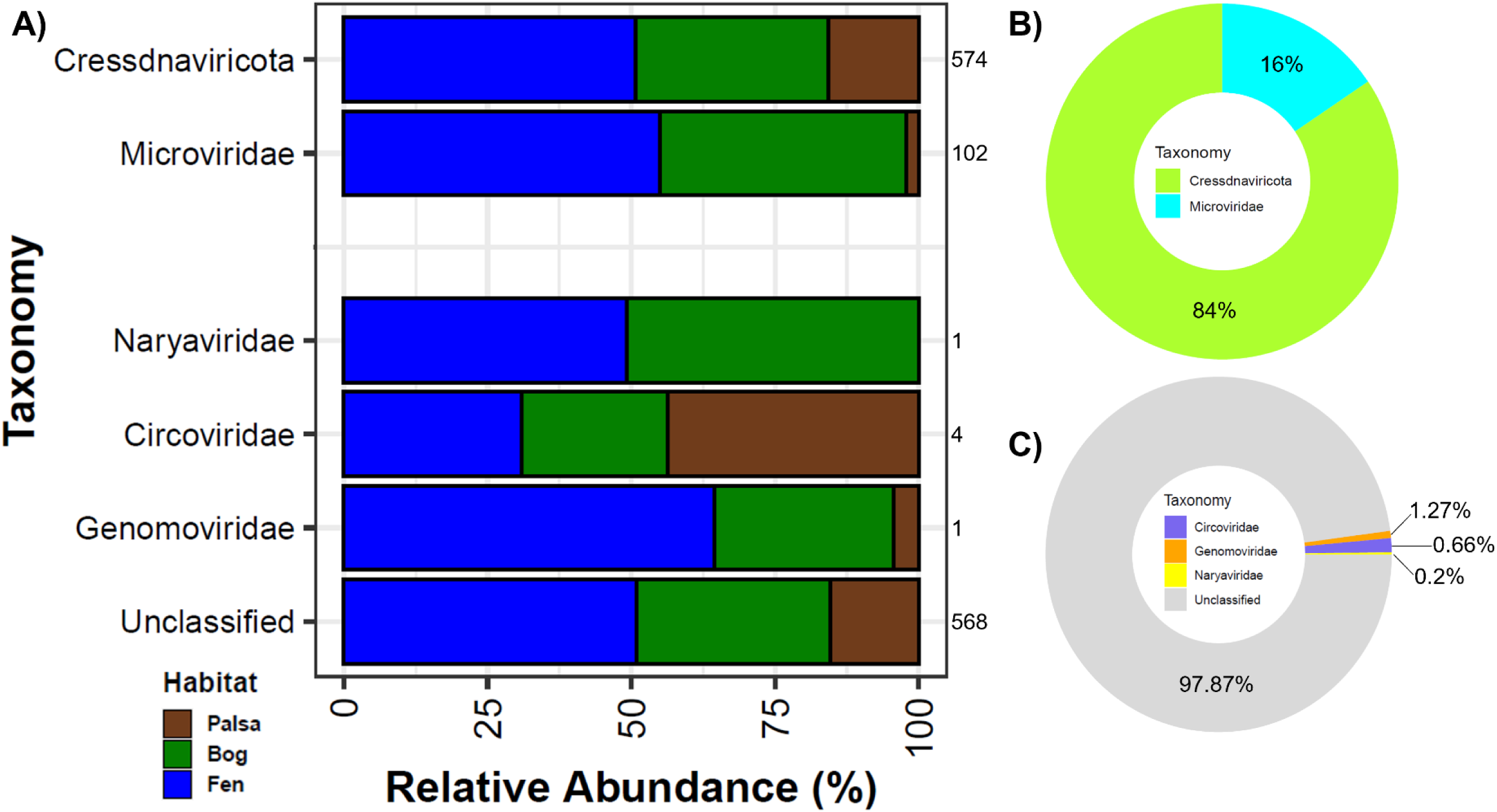
Taxonomy and Ecology of recovered ssDNA vOTUs across Stordalen Mire. (A) A stacked bar chart showing the coverage of the ssDNA vOTUs taxonomically across the habitats. The number at the top of each bar represents the number of vOTUs detected. (B) A Pie chart showing the coverage of ssDNA vOTUs from the family *Microviridae* and the phylum *Cressdnaviricota*, and (C) a further taxonomic breakdown of *Cressdnaviricota* coverage at the family level.

The coverage of the ssDNA vOTUs was used to assess their distribution along the permafrost thaw gradient. The ssDNA vOTUs were most abundant in the fen samples, similar to the dsDNA vOTUs, accounting for more than half of the coverage (Fig. 3C). This result holds true for both types of ssDNA viruses detected, but the ssDNA vOTU signal was dominated by cressdnaviruses (86%) and one fen sample (51%, from sample F_XD_2016; Supplementary Table 7). When we compare the ssDNA vOTUs between the years and depths sampled (i.e., 2016 to 2017 and middle and extra deep samples), the richness was 1.6-fold higher for cressdnaviruses and 10-fold higher for microviruses in 2016 compared to 2017, and interestingly the cressdnaviruses switched from being most abundant in the extra deep sample in 2016 to the middle depth sample in 2017. These results are dramatically different compared to a previous characterization of ssDNA vOTUs from this field site (Trubl et al. 2019), where viruses in the family *Microviridae* were not detected in the palsa habitat and the fen habitat had the lowest viral richness, suggesting more spatiotemporal sampling at smaller/shorter intervals may be needed to resolve these viral population dynamics.

### Global comparison of vOTUs

In recent years, large-scale viromics has been performed on marine, human gut, and soil ecosystems, therefore we compared these datasets, along with viral isolate genomes and previous soil virus datasets from Stordalen Mire, to assess the global distribution of our vOTUs (Supplementary Table 4). Not surprisingly ∼97.5% of the vOTUs from the combined global dataset included exclusively environmental sequences (i.e., metagenomes and viromes) and not isolates, and >95% from viromes, highlighting the capability of these targeted metagenomes to capture more vOTUs (which may not be true for the human gut, see Gregory et al. 2020). Also not surprising, a comparison of vOTUs derived from marine, human gut, and soil systems revealed very little overlap, with >99.9% of vOTUs specific to only one of these vastly different biomes (Fig. 4A). This confirms previous assessments of habitat or ecological zone specificity for viruses where vOTUs and viral communities are structured by environmental conditions (Brum et al. 2015; Trubl et al. 2018; Emerson et al. 2018; Gregory et al. 2019; ter Horst et al. 2021; Lee et al. 2022b). Shifting from completely global to more soil-focused comparisons, we next explored how these new viruses compared to previously collected viruses from Stordalen Mire and other soils. Analogously, most (>97%) of our vOTUs were novel. Of the 3% of vOTUs that were shared, the vast majority (224 of 273) were seen in previous Stordalen Mire datasets (SMV1: Trubl et al. 2018; Trubl et al. 2019 and SMM: Emerson et al. 2018) versus 9 shared with other soils, and 5 with RefSeq (Fig. 4B). There were no vOTUs shared with agricultural soils (Santos-Medellin et al. 2021) or biocrusts (Van Goethem et al. 2019), nor with permafrost-associated bog soils from Alaska (ICE-SIP: Trubl et al. 2021), although those soils did have one vOTU shared with a previous Stordalen Mire viromic dataset (SMV1). The other peatland dataset (SPRUCE: Ter Horst et al. 2021) did have three shared vOTUs with our dataset (SMV2) and eight vOTUs shared with previous Stordalen Mire datasets (SMV1 and SMM). Altogether there is high genetic novelty in each of the soil datasets, especially considering the small amount of shared vOTUs in our dataset compared to previous Stordalen Mire datasets. The minimal overlap in shared vOTUs among the Stordalen Mire datasets is presumably due to differences in virome vs bulk metagenome sampling strategies (i.e., metagenomes in SMM with 9% shared and viromes in SMV1 with 24% shared).

**Figure 4.**
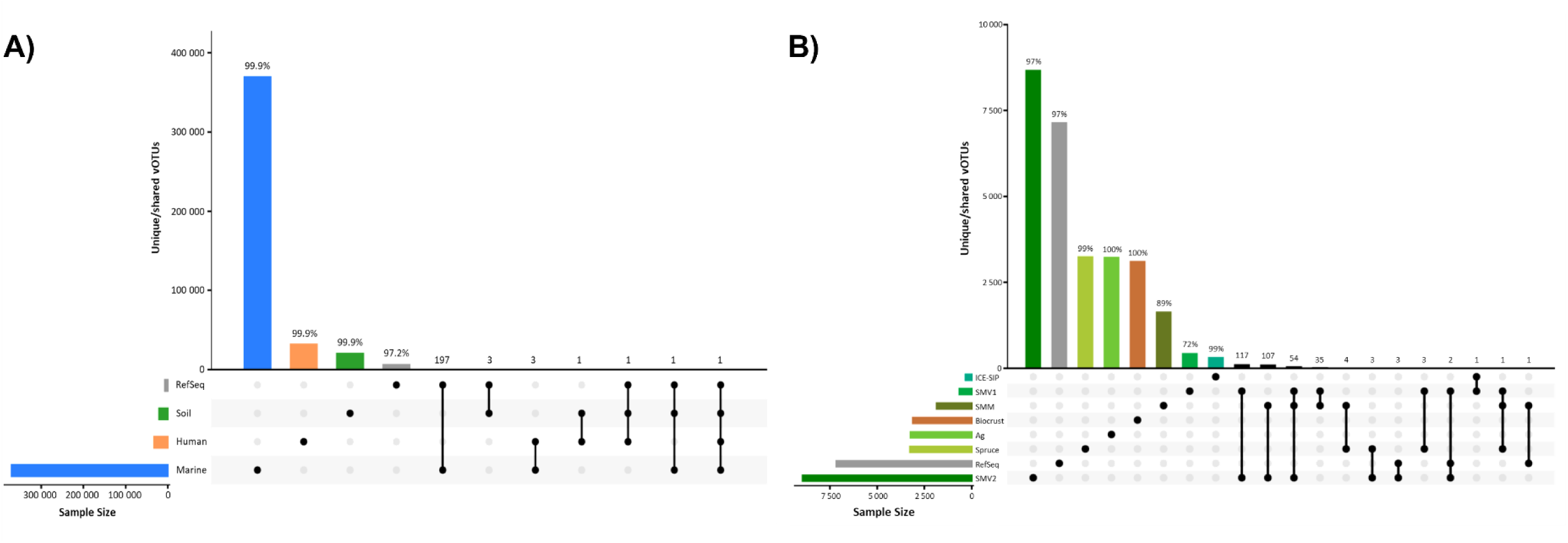
Global comparison of Stordalen Mire dsDNA vOTUs to marine, human and soil viruses. (A) Comparison of unique and shared viral populations (vOTUs) among marine, human, soil, and RefSeq (v204) datasets and (B) a comparison only among soil datasets. The columns represent the percent of unique vOTUs (left side of each figure) or the number of shared vOTUs per dataset (right side of the figure), with a line below the column denoting the datasets being compared. The rows represent each dataset. (A) Datasets are color coded by habitat with blue for marine, orange for human, green for soil, and gray for RefSeq. (B) Datasets are color coded by source and gray for RefSeq. Datasets: GOV2 (marine; viromes; Gregory et al. 2019), GVD (human gut; metagenomes and viromes; Gregory et al. 2020), SMV2 (soil; viromes; this paper), SMV1 (soil; viromes; Trubl et al. 2018, Trubl et al. 2019), SPRUCE (soil; viromes; ter Horst et al. 2021), Ag (soil; viromes; Santos-Medellin et al. 2020), Biocrust (soil; metagenomes; Van Goethem et al. 2019), SMM (soil; metagenomes, viromes removed; Emerson et al. 2018), ICE SIP (soil; stable isotope probing metagenomes; Trubl et al. 2021).

Although there has been significant activity in identifying ssDNA viruses from environmental systems (López-Bueno et al. 2009; Zawar-Reza et al. 2014; Aguirre de Cárcer et al. 2015; Dayaram et al. 2016; Sommers et al. 2019; de La Higuera et al. 2020), there is limited work on soils which makes a global comparison challenging. We took advantage of the two marker genes (MCP and Rep) used to capture ssDNA viruses to further explore their genetic novelty and compared ssDNA viruses from GenBank (Oct. 5, 2021) and the Les Pradeaux Mire (a sphagnum-dominated peatland in France) (Quaiser et al. 2015; 2016). Only four microviruses from our dataset clustered with those of identified in Les Pradeaux Mire (Fig. 5A) (Quaiser, 2015) and 29 cressdnaviruses phylogenetically clustered with 15 (of the 34) Rep proteins from ssDNA viruses identified in Les Pradeaux Mire (Fig. 5B) (Quaiser et al. 2016). The minimal overlap of ssDNA viruses from Stordalen Mire with another peatland indicates there is still a lot of novel ssDNA virus sequence space in soils, undoubtedly the result of high mutation rates of ssDNA viruses and that microviruses may be the most abundant ssDNA viruses on the planet (Kirchberger et al. 2022).

**Figure 5.**
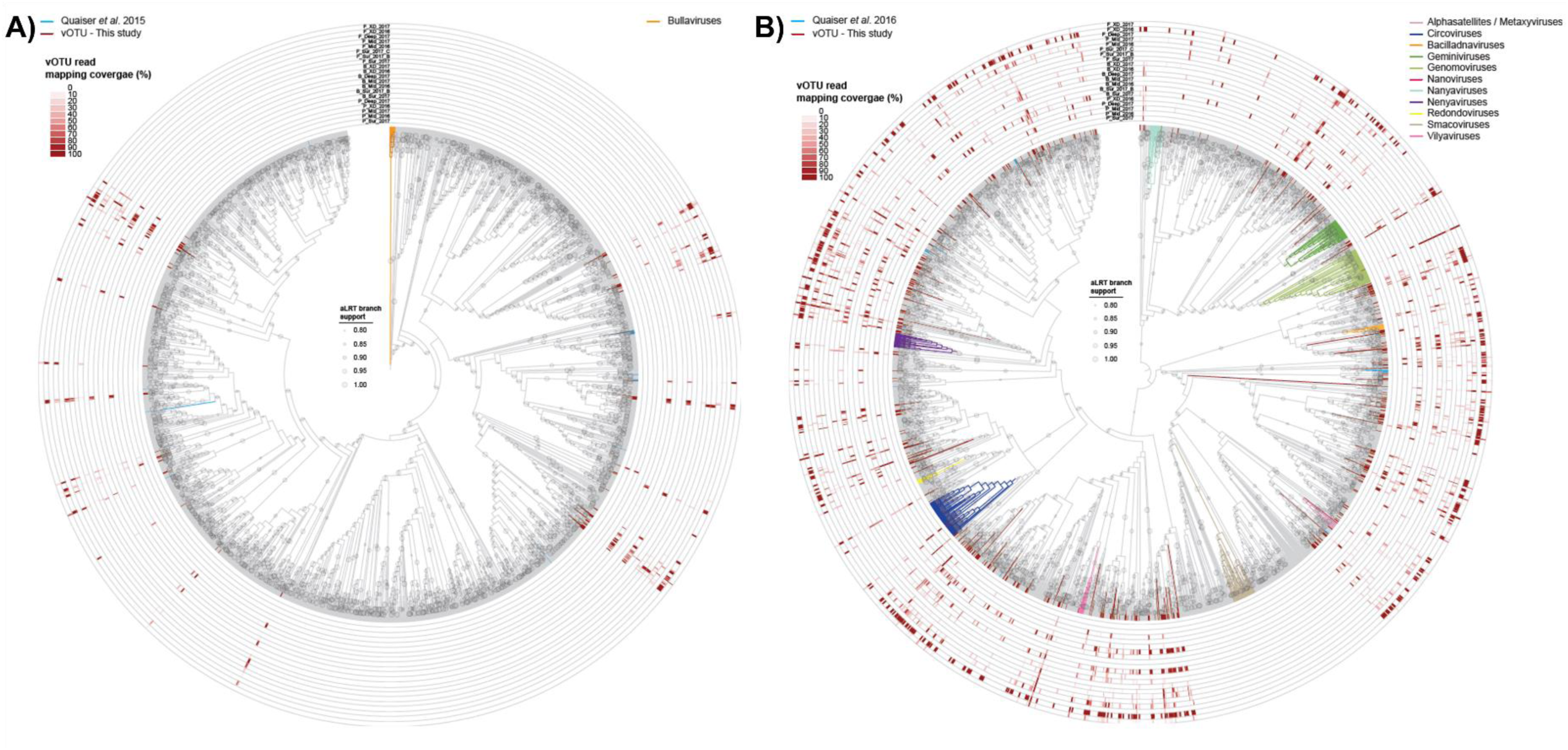
Phylogenetic comparisons of ssDNA vOTUs to Les Pradeaux Mire. Maximum likelihood phylogenies of (A) MCP amino acid sequences encoded by the microvirus genomes or (B) Rep amino acid sequences encoded by cressdnaviruses available in GenBank as well as those representing vOTUs from this study. The phylogenies are represented as cladograms and aLRT branch support >0.8 is shown. The MCP phylogenetic tree is rooted with sequences of bullaviruses represented with an orange clade and MCPs are shown in red nodes. The Rep phylogenetic tree is midpoint rooted and classified cressdnavirus Reps in the phylogeny are shown with color coded clades whereas those from this study are shown in red nodes. In both phylogenies the representatives from the *sphagnum*-dominated peatland Les Pradeaux Mire (MCP from Quaiser et al. 2015; Rep from Quaiser et al. 2016) are shown in blue. The percentage genome coverage of the mapped reads from each sample to the vOTUs is shown on the outside of the cladograms.

### Inferring possible ecosystem impacts of soil viruses

We next sought to predict hosts for this study’s 9,560 vOTUs to better understand their impacts on microbial biogeochemistry. We first used CRISPR spacers to match vOTUs to 1,529 MAGs previously reconstructed from Stordalen Mire (Woodcroft et al. 2018). This resulted in extraction of 3,560 spacer sequences from CRISPRs identified in the 1,529 MAGs and there were 42 spacers matching our vOTUs, with 38 unique spacers (from 23 host CRISPRs) matching to 13 unique dsDNA vOTUs to 24 unique MAGs (Supplementary Table 5). Even with the four-to-six years’ difference in sampling (2010-2012 for the MAGs, 2016-2017 for this study’s vOTUs), the number of CRISPR spacers from a host that matched a vOTU ranged from one to five spacers, demonstrating the utility of CRISPR spacers to link viruses and microbial hosts even when not sampled simultaneously. We detected 36 unique links between dsDNA vOTUs and MAGs from phyla Verrucomicrobiota and Acidobacteriota, with roughly three quarters to MAGs from Verrucomicrobiota subdivision 3 (26 of 36), one quarter to Acidobacteriotaceae MAGs (9 of 36), and 1 match to a Solibacterales MAG. Beyond CRISPR-based host predictions, we looked next at using nucleotide similarity matches to target potentially integrated viruses and identified 644 novel putative hosts for 488 vOTUs (i.e., there was no overlap between host linkage methods; Supplementary Table 5) and expanding the number of linked microbial hosts to 12 bacterial phyla and 1 archaeal phylum (Fig. 6). The most “infected” phylum of bacteria was Acidobacteriota, accounting for more than half of the total virus-host linkages (53%), followed by Actinobacteria (15%), Verrucominbacteria (11%), and Patescibacteria (11%). Acidobacteriota is the most abundant phylum in the bog (Woodcroft et al 2018) where viral richness and diversity were highest, and its dominance of virus-host linkages was seen in previous Stordalen Mire virus-host surveys (Emerson et al 2018; Trubl et al. 2018).

**Figure 6.**
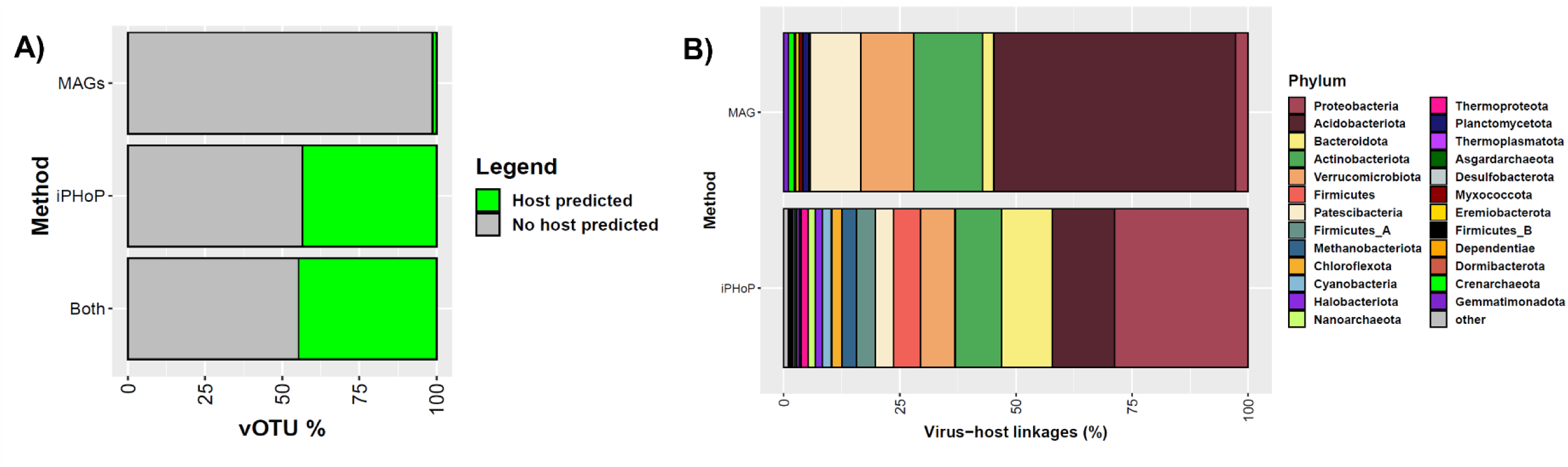
Virus-host linkages along the permafrost thaw gradient. (A) A stacked bar chart showing the number of vOTUs with predicted hosts from the Stordalen Mire MAGs (126; 1.3%), iPHoP with the GTDB database (4,149; 43.4%), and the total number of vOTUs with host linkages (4,275; 44.7%). (B) A stacked bar chart showing the top 25 microbial host phyla predicted from Stordalen Mire MAGs previously recovered (Woodcroft et al. 2018) or iPHoP with the GTDB database (release 202). The “other” category represents a summation of the remaining virus-host linkages (see Supplementary Table 5). For iPHoP, confidence in taxonomic rank is dependent on score and available host resolution in GTDB database (see methods).

Host prediction of viruses is an arduous process with our effort only linking 1.3% of our vOTUs to MAGs, and to be able to maximize the number of putative hosts for our vOTUs, we next compared our dsDNA vOTUs to the GTDB database (release 202) using the iPHoP bioinformatic tool trained on the 1,529 MAGs from Stordalen Mire (see Methods). In total, there were 7,815 virus-host linkages obtained with iPHoP, in which we predicted hosts for 4,149 vOTUs (43%) spanning bacteria (92%) and archaea (8%), and 49 phyla; including 1,150 vOTUs with scores ≥90 allowing genus-level taxonomic resolution for the predicted host (Supplementary Table 5). This global prediction effort increased our host predictions by almost 9-fold and yielded similar predicted hosts at the phylum level (top phyla: Proteobacteria, Bacteroidota, Actinobacteriota, Acidobacteriota, Verrucomicrobiota) but a different host community at lower taxonomic ranks (top shared genera: UBA11358, Terracidiphilus; Fig. 6).

Current host prediction tools for viruses were developed using dsDNA viruses, so we predicted hosts for ssDNA vOTUs based on similar nucleotide content to known ssDNA virus hosts. Based on our phylogenic comparison of microviruses with those identified in another peatland, Les Pradeaux Mire (Fig. 4C) (Quaiser, 2015), we cautiously infer four microviruses vOTUs to infect bacteria from the phylum Bacteroidota. *Microviridae* is a family of ssDNA bacteriophage with a wide host range that mainly infect *Enterobacteria* (intracellular parasitic Gram-negative bacteria), *Spiroplasma* (parasitic bacteria without a cell wall), and *Rhizobiaceae* (Gram-negative nitrogen fixing bacteria that have symbiotic relationship with plants and often dominate *Sphagnum* peatlands), and to a lesser extent other Gram-negative bacteria (Brentlinger et al. 2002; Krupovic and Forterre 2011; Kolton et al. 2022; Kirchberger et al. 2022). This suggests that although microviruses may not carry AMGs nor participate in horizontal gene transfer (Kirchberger et al. 2022), they may still have appreciable impacts on the soil ecosystems. Particularly in fen ecosystems their putative host, Bacteriodota, are the primary lactate metabolizers in the fen (Woodcroft et al. 2018) and have the capability to degrade a wide range of complex carbohydrates via their polysaccharide utilization loci (McKee et al. 2021).

Cressdnaviruses are known to infect a wide array of eukaryotes (Zhao, Lavington, Duffy 2021) and twelve of the cressdnavirus vOTUs have identifiable ORFs that are Rep proteins when using a ciliate translation table, which we interpret to suggest that they infect ciliates here too. Ciliates are bacterivorous protozoan that are commonly found in the active layer of permafrost soils, and along with other protozoa play an important role in controlling bacterial assemblages and recycling soil organic matter (Coolen et al. 2011; Schostag et al. 2019). The roles of ciliates may increase as permafrost thaw continues and these environments become inundated with water, suggesting their viruses will also become more abundant and play larger roles in C cycling. Additionally, one of the cressdnavirus vOTUs fell into the newly created genus *Naryaviridae*, which are only known to infect Entamoeba (Krupovic and Varsani, 2022). Entamoeba is a genus of Amoebozoa, they are internal parasites of animals, and are known to be able to survive in permafrost (Shmakova and Rivkina, 2015). These protists can be infected by ssDNA viruses (Barreat and Katzourakis, 2022.), giant DNA viruses (Aherfi et al. 2016), and RNA viruses (Wu et al. 2022), and may provide a means for viruses to persist in permafrost.

### Stordalen Mire viruses are associated with dominant and major C cycling microbial lineages

Beyond infecting key C cycling microbes in Stordalen Mire, we next explored whether the viruses might impact ecosystem biogeochemistry by carrying AMGs or metabolic genes that may augment host microbial metabolism. The dsDNA vOTUs were investigated using DRAM-v (Shaffer et al. 2020) with an additional, customized database reflecting carbohydrate utilization profiles of the MAGs from the same system (Supplementary Table 6; Woodcroft et al. 2018). DRAM-v identified 1,648 putative AMGs or metabolic genes from 959 vOTUs with a score of 1–3. Of the 1,648 putative AMGs and metabolic genes, 644 were novel, and the other 994 were previously reported, 46 of which were previously experimentally verified, i.e., shown in a microbial host to provide a specific function (Fig. 7A). Functions of the genes were assigned to C utilization (47%), organic nitrogen metabolism (12%), miscellaneous (39%), energy metabolism (2%), and transporter (∼1%) DRAM-v categories (Fig. 7B). In each category, certain subcategories were more prevalent, such as Carbohydrate Active Enzymes (CAZymes) in C utilization, amino acid utilization and transformations in organic nitrogen, information systems in miscellaneous, and sulfur metabolism in energy. Interrogation of the novel AMGs and metabolic genes found they spanned all five DRAM-v categories, representing 102 novel functions, and notably half (51%) were CAZymes with 59 different functions. The detection of AMGs and the diversity in AMG functions increased with thaw (22% identified in virus genomes from palsa and ∼40% in bog and fen; Supplementary Table 6), mirroring microbial and plant compound diversity (Woodcroft et al. 2018; Hough et al. 2022). We leveraged our virus-host linkages to provide the lineage-specific microbial metabolism that these AMGs and metabolic genes may be augmenting and identified 19 dsDNA vOTUs linked to bacteria in the Stordalen Mire MAGs that carried 75 genes.

**Figure 7.**
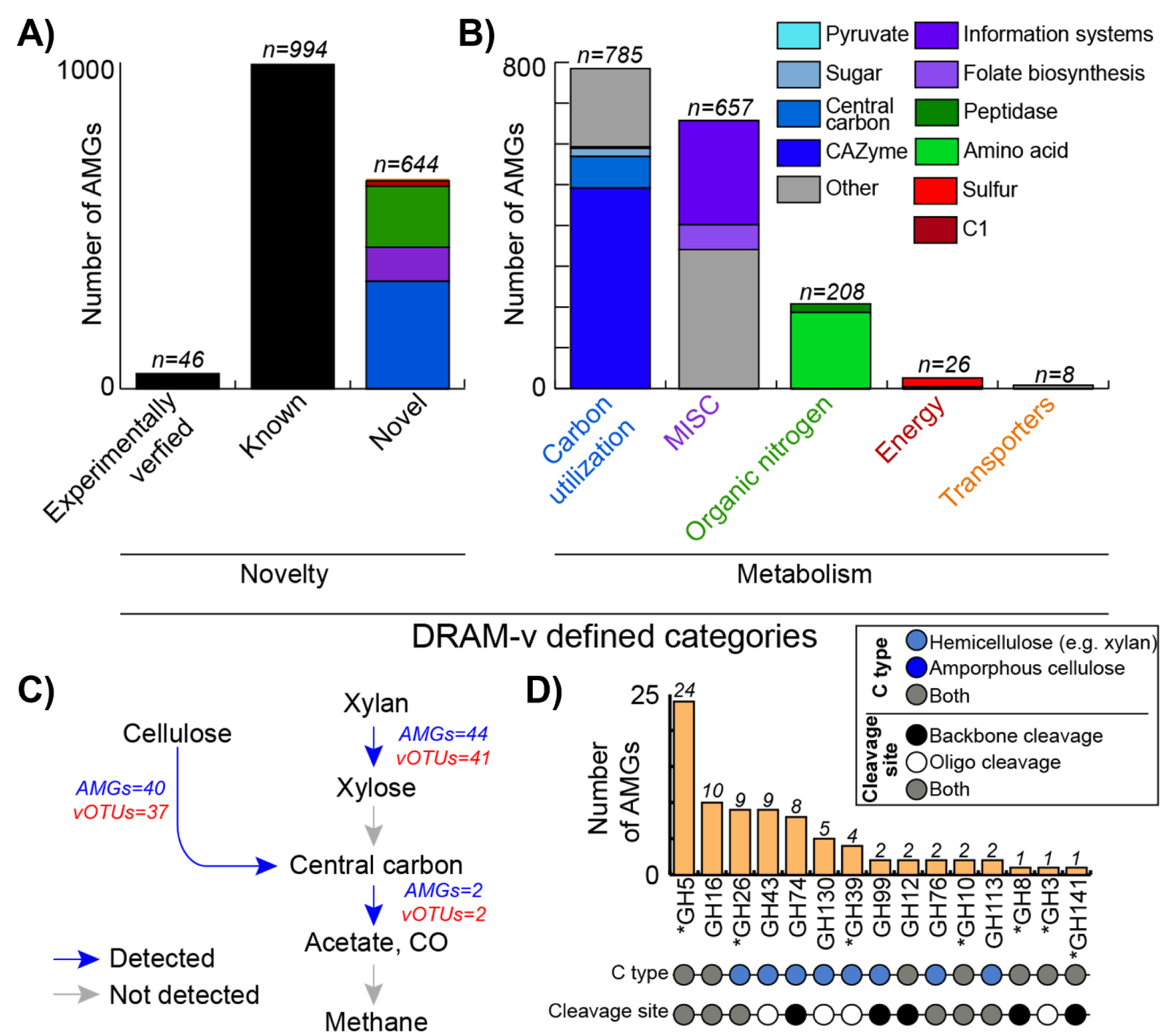
Putative AMGs and metabolic genes guided by dominant microbial metabolisms. (A) The portion of the 1,648 putative AMGs recovered by DRAM-v that have been experimentally verified, previously reported (known), and previously unreported (novel) 1,648 separated into the five major functional categories. (B) The metabolic assignments of the 1,684 AMGs, separated into the five major functional categories and subcategories. (C) Cellulose and xylan (hemicellulose) degradation pathways feeding into methanogenesis as reported in Woodcroft et al. (2018). Blue arrows denote genes in the pathways were detected in viral genomes, and gray denotes no genes were detected. For each pathway step, the number of AMGs is shown in blue text, the number of vOTUs encoding those AMGs is in red text. (D) The distribution of the 15 families of glycoside hydrolases (GH) carried by vOTUs that can degrade cellulose or hemicellulose, and the number of GHs detected. The asterisks denote those families that can use xylan. The dot plot specifies the GH’s function and the cleavage site.

Focusing on the AMGs involved in C cycling, which was the most abundant metabolic gene type detected, these genes were involved in many important facets of microbial C cycling previously identified in Stordalen Mire (Woodcroft et al. 2018), including degradation of large C polymers (polysaccharides) and C monomers (monosaccharides) for utilization of sugars, and short chain fatty acid interconversions that can supply the substrates needed for methanogenesis (Fig. 7C). Previous Stordalen Mire research showed that more than half of the microbial populations that had the potential to degrade cellulose also had the capacity for xylan degradation, with most xylan degradation activity (inferred by metatranscriptomics) occurring by Actinobacteriota in the palsa, Acidobacteriota, Actinobacteriota, and Verrucomicrobiota in the bog, and Bacteroidota, mainly Ignavibacteria, in the fen. Most of the identified microbial hosts belonged to the phyla Acidobacteriota, Actinobacteriota, Bacteroidota, and Verrucomicrobiota. It is likely that Acidobacteriota (cellulase- and xylanase-encoding) are the primary degraders of large polysaccharides in the palsa and bog (61% and 75% of Acidobacteriota genomes, respectively; Woodcroft et al. 2018.), revealing the high potential for viruses to dictate the fate of the complex C that makes up most of the peat matrix (Hodgkins et al. 2014). Interestingly, most Acidobacteriota populations did not have the capability for xylan degradation, but did encode genes for xylose degradation, and in our data, we found 44 AMGs for xylan degradation carried by 41 vOTUs, but none for xylose degradation (Fig. 7C). While none of our virus-host linkages could provide information on which microbes these viruses infect, we posit viruses carry these AMGs alleviating a bottleneck in this microbial degradation pipeline contributing to the formation of more labile products promoting C loss from these peatlands.

We continued our investigation of viral roles in the microbial degradation pipeline starting with complex carbohydrate degradation and continuing down to degradation of more labile compounds. The most prevalent C cycling AMGs and metabolic genes encoded for CAZymes, with 278 vOTUs carrying 345 glycosyltransferases (spanning 31 types), which catalyze the formation of glycosidic bonds and 112 vOTUs carrying 123 glycoside hydrolases (GHs, spanning 32 types), which catalyze the hydrolysis of glycosidic bonds. Focusing on the GHs that enable the turnover of complex carbohydrates to smaller more labile C compounds, the most abundant GHs belonged to GH5 (n=24), GH16 (n=10), GH26 (n=9) and GH43 (n=9) families (Fig. 7D). Notably, each of these GHs have the capacity to degrade cellulose or hemicellose (e.g., xylan), both of which were important polysaccharide substrates for microorganisms in Stordalen Mire. Once these substrates are degraded, they can be fuel for other microbial metabolisms and recently, these metabolisms were identified to be prevalent and play a major role in Stordalen Mire CH_4_ dynamics (Woodcroft et al. 2018), which viral abundance was found to help predict (Emerson et al. 2018). We identified 53 vOTUs that carried genes for the degradation of monosaccharides such as mannose (via mannose-6-phosphate isomerase or mannose 4,6-dehydratase) and galactose (via UDP-glucose 4-epimerase or glucose dehydrogenase), and two of the vOTUs contained seven AMGs that were linked to hosts from Acidobacteriota. At lower trophic levels, AMGs were identified with the potential for short chain fatty acid and alcohol interconversions, including acetate kinase (n=1), acylphosphatase (n=1), and acetate dehydrogenase (n=1) aiding in reduction of C1 compounds; short chain fatty acids could be used by acetoclastic methanogens to produce CH_4_. Acetate could also be scavenged by Firmicutes (which account for 0.5% of the 1,529 MAGs) and used to produce butyrate or propionate (Mahowald et al. 2009; Tan et al. 2014). If the latter is produced, then Deltaproteobacteria (6.2% of MAGs) can convert the propionate to acetate, syntrophically fueling acetoclastic methanogenesis (Wawrik et al. 2016), which has been found previously in peatlands (Schmidt et al. 2016) and was previously suggested for Stordalen Mire (Trubl et al. 2018). Stordalen Mire is a peatland and is currently thought to store C, but with permafrost thaw it is becoming a productive wetland, releasing C as CO_2_ and CH_4_ through processes primarily driven by microbes (Woodcroft et al. 2018). Identification of Stordalen Mire viruses that carry C cycling AMGs and metabolic genes, and linkages to known dominant hosts with C-cycling lifestyles, suggests these soil viruses are a fulcrum in soil C cycling.

## Conclusions

Climate change is driving permafrost thaw in these peatlands and causing a shift from low productivity palsa to more productive fen habitats which is increasing plant compound and microbial diversity. Although these peatlands are currently C sinks, this feature depends on the organic matter content available and the microbial inhabitants. In this study, we characterized thousands of novel ssDNA and dsDNA virus populations and show that viral diversity increases with thaw and the functional diversity of the genes viruses carry to augment host metabolism. These viruses likely affect C cycling by participating in biomass recycling of complex polysaccharides, revealing the ecological roles of viruses in these climate critical ecosystems. The contextualization of Stordalen Mire viruses revealed additional mechanistic insights into microbial metabolic reprogramming that affects genome-scale metabolic modeling, putting an increased emphasis on the need to better contextualize viruses in soils and in biogeochemical models. Given the global relevance of peatlands and permafrost in the C cycle and the detection of C cycling genes in indigenous viruses, the role of these viruses in global C cycle could be significant.

## Acknowledgements

We thank Moira Hough, Sky Dominguez, and Nicole Raab for collecting soil cores and geochemical data, and the Abisko Naturvetenskapliga Station for field support. This study was funded by the Genomic Science Program of the U.S. Department of Energy (DOE) Office of Biological and Environmental Research (BER) (grants DE-SC0010580, DE-SC0016440, DE-SC0248445, DE-SC002330), a Gordon and Betty Moore Foundation Investigator Award (GBMF#3790), and by the National Science Foundation, Biology Integration Institutes Program Award (#2022070). G.T. was supported by a U.S. DOE Office of Science Graduate Student Research (SCGSR) program award, a U.S. DOE, Office of BER, Genomic Science Program ‘Microbes Persist’ Scientific Focus Area award (#SCW1632), and a U.S. DOE Laboratory Directed Research & Development award (21-LW-060). Work conducted at Lawrence Livermore National Laboratory was under the auspices of the U.S. DOE under contract DE-AC52-07NA27344. The SCGSR program is administered by the Oak Ridge Institute for Science and Education and is managed by ORAU (contract number DE-SC0014664). The work (proposal: 10.46936/10.25585/60000758) conducted by the U.S. DOE Joint Genome Institute (https://ror.org/04xm1d337), a DOE Office of Science User Facility, is supported by the Office of Science of the U.S. DOE operated under Contract No. DE-AC02-05CH11231.

---

**Supplementary Table 1. Geochemistry of samples.**

**Supplementary Table 2. Sample preparation and sequencing information.**

**Supplementary Table 3. List of vOTUs and genomic information.**

**Supplementary Table 4. Global dataset information.**

**Supplementary Table 5. Information on virus-host linkages via MAGs and iPHoP.**

**Supplementary Table 6. List of AMGs and other metabolic genes identified via DRAM-v.**

**Supplementary Table 7. Coverage profile of ssDNA vOTUs.**

**Supplementary Figure 1. Viral contig recovery and coverage thresholds of viruses from each library method.** (A) A comparison of viral contigs recovered for each sample from each library method. Comparisons of normalized coverage for each sample at the relaxed (B) or strict (C) threshold. Comparisons of normalized coverage for each viral contig recovered in both library kits at the relaxed (D) or strict (E) threshold. The three outliers (see methods) are indicated in orange and are based on contig recovery.

**Supplementary Figure 2.** (A) A heatmap comparing samples grouped by habitat and depth and (B) a heatmap of the 20 samples clustered using a Bray-Curtis dissimilarity metric based on vOTU normalized abundance.

## References

Aguirre de Cárcer, D., López-Bueno, A., Pearce, D. A., & Alcamí, A. (2015). Biodiversity and distribution of polar freshwater DNA viruses. Science advances, 1(5), e1400127.

Aherfi, S., Colson, P., La Scola, B., & Raoult, D. (2016). Giant viruses of amoebas: an update. Frontiers in microbiology, 7, 349.

Åkerman, H. J., & Johansson, M. (2008). Thawing permafrost and thicker active layers in sub-arctic Sweden. Permafrost and periglacial processes, 19(3), 279–292.

Altschul, S. F., Gish, W., Miller, W., Myers, E. W., & Lipman, D. J. (1990). Basic local alignment search tool. Journal of Molecular Biology, 215(3), 403–410. doi:10.1016/s0022-2836(05)80360-2

Barnett, S. E., & Buckley, D. H. (2023). Metagenomic stable isotope probing reveals bacteriophage participation in soil carbon cycling. Environmental Microbiology.

Barreat, J. G. N., & Katzourakis, A. (2022). Paleovirology of the DNA viruses of eukaryotes. Trends in Microbiology, 30(3), 281–292.

Bland, C., Ramsey, T. L., Sabree, F., Lowe, M., Brown, K., Kyrpides, N. C., & Hugenholtz, P. (2007). CRISPR Recognition Tool (CRT): a tool for automatic detection of clustered regularly interspaced palindromic repeats. BMC Bioinformatics, 8(1), 209. doi:10.1186/1471-2105-8-209

Bolduc, B., Hodgkins, S. B., Varner, R. K., Crill, P. M., Mccalley, C. K., Chanton, J. P., Tyson, G. W., Riley, W. J., Palace, M., Duhaime, M. B., Hough, M. A., IsoGenie Project Coordinators, IsoGenie Project Team, A2A Project Team, Saleska, S. R., Sullivan, M. B., Rich V. I. (2020). The IsoGenie database: an interdisciplinary data management solution for ecosystems biology and environmental research. PeerJ, 8, e9467. doi:10.7717/peerj.9467

Breitbart, M. (2012). Marine viruses: truth or dare. Annual review of marine science, 4(1), 425–448.

Brentlinger, K. L., Hafenstein, S., Novak, C. R., Fane, B. A., Borgon, R., McKenna, R., & Agbandje-McKenna, M. (2002). Microviridae, a family divided: isolation, characterization, and genome sequence of φMH2K, a bacteriophage of the obligate intracellular parasitic bacterium Bdellovibrio bacteriovorus. Journal of bacteriology, 184(4), 1089–1094.

Brettin, T., Davis, J. J., Disz, T., Edwards, R. A., Gerdes, S., Olsen, G. J., Olson, R., Overbeek, R., Parrello, B., Pusch, G. D., Shukla, M., Thomason III, J. A., Stevens, R., Vonstein, V., Wattam, A. R., & Xia, F. (2015). RASTtk: a modular and extensible implementation of the RAST algorithm for building custom annotation pipelines and annotating batches of genomes. Scientific reports, 5(1), 8365.

Bruinsma, S., Burgess, J., Schlingman, D., Czyz, A., Morrell, N., Ballenger, C., Meinholz, H., Brady, L., Khanna, A., Freeberg, L., Jackson, R. G., Mathonet, P., Verity, S. C., Slatter, A. F., Golshani, R., Grunenwald, H., Schroth, G. P., & Gormley, N. A. (2018). Bead-linked transposomes enable a normalization-free workflow for NGS library preparation. BMC Genomics, 19(1), 1–16.

Brum, J. R., Ignacio-Espinoza, J. C., Roux, S., Doulcier, G., Acinas, S. G., Alberti, A., Chaffron, S., Cruaud, C., De Vargas, C., Gasol, J. M., Gorsky, G., Gregory, A. C., Guidi, L., Hingamp, P., Iudicone, D., Not, F., Ogata, H., Pesant, S., Poulos, B. T., Schwenck, S. M., Speich, S., Dimier, C., Kandels-Lewis, S., Picheral, M., Searson, S., Tara Oceans Coordinators, Bork, P., Bowler, C., Sunagawa, S., Wincker, P., Karsenti, E., & Sullivan, M. B. (2015). Patterns and ecological drivers of ocean viral communities. Science, 348(6237), 1261498.

Capella-Gutierrez, S., Silla-Martinez, J. M., & Gabaldon, T. (2009). trimAl: a tool for automated alignment trimming in large-scale phylogenetic analyses. Bioinformatics, 25(15), 1972–1973. doi:10.1093/bioinformatics/btp348

Conway, J. R., Lex, A., & Gehlenborg, N. (2017). UpSetR: an R package for the visualization of intersecting sets and their properties. Bioinformatics, 33(18), 2938–2940. doi:10.1093/bioinformatics/btx364

Coolen, M. J. L., Van De Giessen, J., Zhu, E. Y., & Wuchter, C. (2011). Bioavailability of soil organic matter and microbial community dynamics upon permafrost thaw. Environmental Microbiology, 13(8), 2299–2314. doi:10.1111/j.1462-2920.2011.02489.x

Crowther, T. W., Todd-Brown, K. E., Rowe, C. W., Wieder, W. R., Carey, J. C., Machmuller, M. B., Snoek, B. L., Fang, S., Zhou, G., Allison, S. D., Blair, J. M., Bridgham, S. D., Burton, A. J., Carrillo, Y., Reich, P. B., Clark, J. S., Classen, A. T., Dijkstra, F. A., Elberling, B., Emmett, B. A., Estiarte, M., Frey, S. D., Guo, J., Harte, J., Jiang, L., Johnson, B. R., Kröel-Dulay, G., Larsen, K. S., Laudon, H., Lavallee, J. M., Luo, Y., Lupascu, M., Ma, L. N., Marhan, S., Michelsen, A., Mohan, J., Niu, S., Pendall, E., Peñuelas, J., Pfeifer-Meister, L., Poll, C., Reinsch, S., Reynolds, L. L., Schmidt, I. K., Sistla, S., Sokol, N. W., Templer, P. H., Treseder, K. K., Welker, J. M., & Bradford, M. A. (2016). Quantifying global soil carbon losses in response to warming. Nature, 540(7631), 104–108.

Dayaram, A., Galatowitsch, M. L., Argüello-Astorga, G. R., van Bysterveldt, K., Kraberger, S., Stainton, D., Harding, J. S., Roumagnac, P., Martin, D. P., Lefeuvre, P., Varsani, A. (2016). Diverse circular replication-associated protein encoding viruses circulating in invertebrates within a lake ecosystem. Infection, Genetics and Evolution, 39, 304–316.

de La Higuera, I., Kasun, G. W., Torrance, E. L., Pratt, A. A., Maluenda, A., Colombet, J., Bisseux, M., Ravet, V., Dayaram, A., Stainton, D., Kraberger, S., Zawar-Reza, P., Goldstien, S., Briskie, J. V., White, R., Taylor, H., Gomez, C., Ainley, D. G., Harding, J. S., Fontenele, R. S., Schreck, J., Ribeiro, S. G., Oswald, S. A., Arnold, J. M., Enault, F., Varsani, A., Stedman, K. M. (2020). Unveiling crucivirus diversity by mining metagenomic data. mBio, 11(5), e01410–01420.

Dominguez-Huerta, G., Zayed, A. A., Wainaina, J. M., Guo, J., Tian, F., Pratama, A. A., Bolduc, B., Mohssen, M., Zablocki, O., Pelletier, E., Delage, E., Alberti, A., Aury, J-M. Carradec, Q., Da Silva, C., Labadie, K., Poulain, J., Tara Oceans Coordinators, Bowler, C., Eveillard, D., Guidi, L., Karsenti, E., Kuhn, J. H., Ogata, H., Wincker, P., Culley, A., Chaffron, S., & Sullivan, M. B. (2022). Diversity and ecological footprint of Global Ocean RNA viruses. Science, 376(6598), 1202–1208.

Duhaime, M. B., Deng, L., Poulos, B. T., & Sullivan, M. B. (2012a). Towards quantitative metagenomics of wild viruses and other ultra-low concentration DNA samples: a rigorous assessment and optimization of the linker amplification method. Environmental Microbiology, 14(9), 2526–2537. doi:10.1111/j.1462-2920.2012.02791.x

Duhaime, M. B., & Sullivan, M. B. (2012b). Ocean viruses: Rigorously evaluating the metagenomic sample-to-sequence pipeline. Virology, 434(2), 181–186. doi:10.1016/j.virol.2012.09.036

Eddy, S. R. (2009). A new generation of homology search tools based on probabilistic inference. In Genome Informatics 2009: Genome Informatics Series Vol. 23 (pp. 205–211).

Edwards, R. A., Mcnair, K., Faust, K., Raes, J., & Dutilh, B. E. (2016). Computational approaches to predict bacteriophage– host relationships. FEMS Microbiology Reviews, 40(2), 258–272. doi:10.1093/femsre/fuv048

Emerson, J. B., Roux, S., Brum, J. R., Bolduc, B., Woodcroft, B. J., Jang, H. B., Singleton, C. M., Solden, L. M., Naas, A. E., Boyd, J. A., Hodgkins, Wilson, R. M., Trubl, G., Li, C., Frolking, S., Pope, P. B., Wrighton, K. C., Crill, P. M., Chanton, J. P., Saleska, S. R., Tyson, G. W., Rich, V., I., Sullivan, M. B. (2018). Host-linked soil viral ecology along a permafrost thaw gradient. Nature Microbiology, 3(8), 870–880. doi:10.1038/s41564-018-0190-y

Fu, L., Niu, B., Zhu, Z., Wu, S., & Li, W. (2012). CD-HIT: accelerated for clustering the next-generation sequencing data. Bioinformatics, 28(23), 3150–3152. doi:10.1093/bioinformatics/bts565

Fuhrman, J. A. (1999). Marine viruses and their biogeochemical and ecological effects. Nature, 399(6736), 541–548.

Goordial, J., Davila, A., Greer, C. W., Cannam, R., DiRuggiero, J., McKay, C. P., & Whyte, L. G. (2017). Comparative activity and functional ecology of permafrost soils and lithic niches in a hyper-arid polar desert. Environmental microbiology, 19(2), 443–458.

Gregory, A. C., Zablocki, O., Zayed, A. A., Howell, A., Bolduc, B., & Sullivan, M. B. (2020). The gut virome database reveals age-dependent patterns of virome diversity in the human gut. Cell host & microbe, 28(5), 724–740. e728.

Gregory, A. C., Zayed, A. A., Conceição-Neto, N., Temperton, B., Bolduc, B., Alberti, A., Ardyna, M., Arkhipova, K., Carmichael, M., Cruaud, C., Dimier, C., Domínguez-Huerta, G., Ferland, J., Kandels, S., Liu, Y., Marec, C., Pesant, S.,- Picheral, M., Pisarev, S., Poulain, J., Tremblay, J-E., Vik, D., Babin, M., Bowler, C., Culley, A. I., de Vargas, C., Dutilh, B. E., Iudicone, D., Karp-Boss, L., Roux, S., Sunagawa, S., Wincker P. & Sullivan, M. B. (2019). Marine DNA viral macro-and microdiversity from pole to pole. Cell, 177(5), 1109–1123. e1114.

Guidi, L., Chaffron, S., Bittner, L., Eveillard, D., Larhlimi, A., Roux, S., Darzi, Y., Audic, S., Berline, L., Brum, J.R., Coelho, L.P., Ignacio Espinoza, J. C., Malviya, S., Sunagawa, S., Dimier, C., Kandels-Lewis, S., Picheral, M., Poulain, J., Searson, S., Tara Oceans Consortium Coordinators, Stemmann, L., Not, F., Hingamp, P., Speich, S., Follows, M., Karp-Boss, L., Boss, E., Ogata, H., Pesant, S., Weissenbach, J., Wincker, P., Acinas, S. G., Bork, P., de Vargas, C., Iudicone, D., Sullivan, M. B., Raes, J., Karsenti, E., Bowler C., & Gorsky, G. (2016). Plankton networks driving carbon export in the oligotrophic ocean. Nature, 532(7600), 465–470.

Gunasekera, S., Abraham, S., Stegger, M., Pang, S., Wang, P., Sahibzada, S., & O’Dea, M. (2021). Evaluating coverage bias in next-generation sequencing of Escherichia coli. PloS one, 16(6), e0253440.

Guo, J., Bolduc, B., Zayed, A. A., Varsani, A., Dominguez-Huerta, G., Delmont, T. O., Pratama, A. A., Gazitúa, M. C., Vik, D., Sullivan, M. B. & Roux, S. (2021). VirSorter2: a multi-classifier, expert-guided approach to detect diverse DNA and RNA viruses. Microbiome, 9(1). doi:10.1186/s40168-020-00990-y

Hoaglin, D. C., & Iglewicz, B. (1987). Fine-tuning some resistant rules for outlier labeling. Journal of the American statistical Association, 82(400), 1147–1149.

Hoaglin, D. C., Iglewicz, B., & Tukey, J. W. (1986). Performance of some resistant rules for outlier labeling. Journal of the American Statistical Association, 81(396), 991–999.

Hough, M., McClure, A., Bolduc, B., Dorrepaal, E., Saleska, S., Klepac-Ceraj, V., & Rich, V. (2020). Biotic and environmental drivers of plant microbiomes across a permafrost thaw gradient. Frontiers in Microbiology, 11, 796.

Hough, M., McCabe, S., Vining, S. R., Pickering Pedersen, E., Wilson, R. M., Lawrence, R., Chang, K. Y., Bohrer, G., IsoGenie Coordinators, Riley, W. J., Crill, P. M., Varner, R. K., Blazewicz, S. J., Dorrepaal, E., Tfaily, M. M., Saleska, S. R., Rich, V. I. (2022). Coupling plant litter quantity to a novel metric for litter quality explains C storage changes in a thawing permafrost peatland. Global change biology, 28(3), 950–968.

Howard-Varona, C., Lindback, M. M., Bastien, G. E., Solonenko, N., Zayed, A. A., Jang, H., Andreopoulos, B., Brewer, H. M., Glavina del Rio, T., Adkins, J. N., Paul, S., Sullivan, M. B. & Duhaime, M. B. (2020). Phage-specific metabolic reprogramming of virocells. The ISME Journal, 14(4), 881–895.

Howe, A. C., Jansson, J. K., Malfatti, S. A., Tringe, S. G., Tiedje, J. M., & Brown, C. T. (2014). Tackling soil diversity with the assembly of large, complex metagenomes. Proceedings of the National Academy of Sciences, 111(13), 4904–4909.

Hultman, J., Waldrop, M. P., Mackelprang, R., David, M. M., McFarland, J., Blazewicz, S. J., Harden, J., Turetsky, M. R., McGuire, A. D., Shah, M. B., VerBerkmoes, N.C., Lee, L. H., Mavrommatis, K. & Jansson, J. K. (2015). Multi-omics of permafrost, active layer and thermokarst bog soil microbiomes. Nature, 521(7551), 208–212.

Jones, M. B., Highlander, S. K., Anderson, E. L., Li, W., Dayrit, M., Klitgord, N., Fabani, M. M., Seguritan, V., Green, J., Pride, D. T., Yooseph, S., Biggs, W., Nelson, K. E., & Venter, J. C. (2015). Library preparation methodology can influence genomic and functional predictions in human microbiome research. Proceedings of the National Academy of Sciences, 112(45), 14024–14029.

Katoh, K., & Standley, D. M. (2013). MAFFT Multiple Sequence Alignment Software Version 7: Improvements in Performance and Usability. Molecular Biology and Evolution, 30(4), 772–780. doi:10.1093/molbev/mst010

Kirchberger, P. C., Martinez, Z. A., & Ochman, H. (2022). Organizing the Global Diversity of Microviruses. mBio, e00588–00522.

Kolton, M., Weston, D. J., Mayali, X., Weber, P. K., McFarlane, K. J., Pett-Ridge, J., Somoza, M. M., Lietard, J., Glass, J. B., Lilleskov, E. A., Shaw, A.J., Tringe, S., Hanson, P. J., & Kostka, J. E. (2022). Defining the Sphagnum Core Microbiome across the North American Continent Reveals a Central Role for Diazotrophic Methanotrophs in the Nitrogen and Carbon Cycles of Boreal Peatland Ecosystems. mBio, 13(1), e03714–03721.

Krupovic, M., & Forterre, P. (2011). Microviridae goes temperate: microvirus-related proviruses reside in the genomes of Bacteroidetes. PloS one, 6(5), e19893.

Krupovic, M., & Varsani, A. (2022). Naryaviridae, Nenyaviridae, and Vilyaviridae: three new families of single-stranded DNA viruses in the phylum Cressdnaviricota. Archives of virology, 1–15.

Krupovic, M., Varsani, A., Kazlauskas, D., Breitbart, M., Delwart, E., Rosario, K., Yutin, N., Wolf, Y. I., Harrach, B., Zerbini, F. M., Dolja, V. V., Kuhn, J. H. & Koonin, E. V. (2020). Cressdnaviricota: a virus phylum unifying seven families of Rep-encoding viruses with single-stranded, circular DNA genomes. Journal of virology, 94(12), e00582–00520.

Langmead, B., & Salzberg, S. L. (2012). Fast gapped-read alignment with Bowtie 2. Nature methods, 9(4), 357–359.

Lee, S., Sieradzki, E. T., Hazard, C., & Nicol, G. W. (2022a). Viruses of soil ammonia oxidising archaea identified using a novel DNA stable isotope probing approach for low GC mol% genomes. bioRxiv.

Lee, S., Sieradzki, E. T., Nicolas, A. M., Walker, R. L., Firestone, M. K., Hazard, C., & Nicol, G. W. (2021). Methane-derived carbon flows into host–virus networks at different trophic levels in soil. Proceedings of the National Academy of Sciences, 118(32), e2105124118.

Lee, S., Sorensen, J. W., Walker, R. L., Emerson, J. B., Nicol, G. W., & Hazard, C. (2022b). Soil pH influences the structure of virus communities at local and global scales. Soil Biology and Biochemistry, 108569.

Letunic, I., & Bork, P. (2021). Interactive Tree Of Life (iTOL) v5: an online tool for phylogenetic tree display and annotation. Nucleic Acids Research, 49(W1), W293–W296. doi:10.1093/nar/gkab301

López-Bueno, A., Tamames, J., Velázquez, D., Moya, A., Quesada, A., & Alcamí, A. (2009). High diversity of the viral community from an Antarctic lake. Science, 326(5954), 858–861.

Mackelprang, R., Saleska, S. R., Jacobsen, C. S., Jansson, J. K., & Taş, N. (2016). Permafrost meta-omics and climate change. Annual review of Earth and planetary sciences, 44(1).

Mahowald, M. A., Rey, F. E., Seedorf, H., Turnbaugh, P. J., Fulton, R. S., Wollam, A., Shah, N., Wang, C., Magrini, V., Wilson, R. K., Cantarel, B.L., Coutinho, P. M., Henrissat, B., Crock, L. W., Russell, A., Verberkmoes, N. C., Hettich, R. L. & Gordon, J. I. (2009). Characterizing a model human gut microbiota composed of members of its two dominant bacterial phyla. Proceedings of the National Academy of Sciences, 106(14), 5859–5864.

McCalley, C. K., Woodcroft, B. J., Hodgkins, S. B., Wehr, R. A., Kim, E. H., Mondav, R., Crill, P. M., Chanton, J. P., Rich, V. I., Tyson, G. W. & Saleska, S. R. (2014). Methane dynamics regulated by microbial community response to permafrost thaw. Nature, 514(7523), 478–481.

McGuire, A. D., Lawrence, D. M., Koven, C., Clein, J. S., Burke, E., Chen, G., Jafarov, E., MacDougall, A. H., Marchenko, S., Nicolsky, D., Peng, S., Rinke, A., Ciais, P., Gouttevin, I., Hayes, D. J., Ji, D., Krinner, G., Moore, J. C., Romanovsky, V., Schädel, C., Schaefer, K., Schuur, E. A. G. & Zhuang, Q. (2018). Dependence of the evolution of carbon dynamics in the northern permafrost region on the trajectory of climate change. Proceedings of the National Academy of Sciences, 115(15), 3882–3887.

McKee, L. S., La Rosa, S. L., Westereng, B., Eijsink, V. G., Pope, P. B., & Larsbrink, J. (2021). Polysaccharide degradation by the Bacteroidetes: mechanisms and nomenclature. Environmental Microbiology Reports, 13(5), 559–581.

Minh, B. Q., Schmidt, H. A., Chernomor, O., Schrempf, D., Woodhams, M. D., Von Haeseler, A., & Lanfear, R. (2020). IQ-TREE 2: New Models and Efficient Methods for Phylogenetic Inference in the Genomic Era. Molecular Biology and Evolution, 37(5), 1530–1534. doi:10.1093/molbev/msaa015

Mondav, R., McCalley, C. K., Hodgkins, S. B., Frolking, S., Saleska, S. R., Rich, V. I., Chanton, J. P. & Crill, P. M. (2017). Microbial network, phylogenetic diversity and community membership in the active layer across a permafrost thaw gradient. Environmental microbiology, 19(8), 3201–3218.

Naccache, S. N., Greninger, A. L., Lee, D., Coffey, L. L., Phan, T., Rein-Weston, A., Aronsohn, A., Hackett Jr, J., Delwart, E. L. &Chiu, C.Y. (2013). The perils of pathogen discovery: origin of a novel parvovirus-like hybrid genome traced to nucleic acid extraction spin columns. Journal of virology, 87(22), 11966–11977.

Nayfach, S., Camargo, A. P., Schulz, F., Eloe-Fadrosh, E., Roux, S., & Kyrpides, N. C. (2020). CheckV assesses the quality and completeness of metagenome-assembled viral genomes. Nature Biotechnology. doi:10.1038/s41587-020-00774-7

Nicolas, A. M., Sieradzki, E. T., Pett-Ridge, J., Banfield, J. F., Taga, M. E., Firestone, M. K., & Blazewicz, S. J. (2022). Isotope-enrichment reveals active viruses follow microbial host dynamics during rewetting of a California grassland soil. bioRxiv.

Nurk, S., Bankevich, A., Antipov, D., Gurevich, A., Korobeynikov, A., Lapidus, A., Prjibelsky, A., Pyshkin, A., Sirotkin, A., Sirotkin, Y., Stepanauskas, R., McLean, J., Lasken, R., Clingenpeel, S. R., Woyke, T., Tesler, G., Alekseyev, M. A. & Pevzner, P. A. (2013). Assembling genomes and mini-metagenomes from highly chimeric reads. In Research in Computational Molecular Biology: 17th Annual International Conference, RECOMB 2013, Beijing, China, April 7-10, 2013. Proceedings 17 (pp. 158–170). Springer Berlin Heidelberg.

Nurk, S., Meleshko, D., Korobeynikov, A., & Pevzner, P. A. (2017). metaSPAdes: a new versatile metagenomic assembler. Genome research, 27(5), 824–834.

Paez-Espino, D., Eloe-Fadrosh, E. A., Pavlopoulos, G. A., Thomas, A. D., Huntemann, M., Mikhailova, N., Rubin, E., Ivanova, N. N. & Kyrpides, N.C., (2016). Uncovering Earth’s virome. Nature, 536(7617), 425–430.

Paez-Espino, D., Pavlopoulos, G. A., Ivanova, N. N., & Kyrpides, N. C. (2017). Nontargeted virus sequence discovery pipeline and virus clustering for metagenomic data. Nature protocols, 12(8), 1673–1682.

Pratama, A. A., Bolduc, B., Zayed, A. A., Zhong, Z. P., Guo, J., Vik, D. R., Gazitúa, M. C., Wainaina, J. M., Roux, S. & Sullivan, M. B., (2021). Expanding standards in viromics: in silico evaluation of dsDNA viral genome identification, classification, and auxiliary metabolic gene curation. PeerJ, 9, e11447.

Quaiser, A., Dufresne, A., Ballaud, F., Roux, S., Zivanovic, Y., Colombet, J., Sime-Ngando, T. & Francez, A.J., (2015). Diversity and comparative genomics of Microviridae in Sphagnum-dominated peatlands. Frontiers in Microbiology, 6, 375.

Quaiser, A., Krupovic, M., Dufresne, A., Francez, A.-J., & Roux, S. (2016). Diversity and comparative genomics of chimeric viruses inSphagnum-dominated peatlands. Virus Evolution, 2(2), vew025. doi:10.1093/ve/vew025

Ren, J., Ahlgren, N. A., Lu, Y. Y., Fuhrman, J. A., & Sun, F. (2017). VirFinder: a novel k-mer based tool for identifying viral sequences from assembled metagenomic data. Microbiome, 5(1). doi:10.1186/s40168-017-0283-5

Rinke, C., Low, S., Woodcroft, B. J., Raina, J. B., Skarshewski, A., Le, X. H., Butler, M. K., Stocker, R., Seymour, J., Tyson, G. W. & Hugenholtz, P. (2016). Validation of picogram-and femtogram-input DNA libraries for microscale metagenomics. PeerJ, 4, e2486.

Roux, S., Adriaenssens, E. M., Dutilh, B. E., Koonin, E. V., Kropinski, A. M., Krupovic, M., Kuhn, J. H., Lavigne, R., Brister, J. R., Varsani, A., Amid, C., Aziz, R. K., Bordenstein, S. R., Bork, P., Breitbart, M., Cochrane, G. R., Daly, R. A., Desnues, C., Duhaime, M. B., Emerson, J. B., Enault, F., Fuhrman, J. A., Hingamp, P., Hugenholtz, P., Hurwitz, B. L., Ivanova, N. N., Labonté, J. M., Lee, K-B., Malmstrom, R. R., Martinez-Garcia, M., Mizrachi, I. K., Ogata, H., Páez-Espino, D., Petit, M-A., Putonti, C., Rattei, T., Reyes, A., Rodriguez-Valera, F., Rosario, K., Schriml, L., Schulz, F., Steward, G. F., Sullivan, M. B., Sunagawa, S., Suttle, C. A., Temperton, B., Tringe, S. G., Thurber, R. V., Webster, N. S., Whiteson, K. L., Wilhelm, S. W., Wommack, K. E., Woyke, T., Wrighton, K. C., Yilmaz, P., Yoshida, T., Young, M. J., Yutin, N., Allen, L. Z., Kyrpides, N. C. & Eloe-Fadrosh, E. A. (2019b). Minimum Information about an Uncultivated Virus Genome (MIUViG). Nature Biotechnology, 37(1), 29–37. doi:10.1038/nbt.4306

Roux, S., Camargo, A. P., Coutinho, F. H., Dabdoub, S. M., Dutilh, B. E., Nayfach, S., & Tritt, A. (2023). iPHoP: An integrated machine learning framework to maximize host prediction for metagenome-derived viruses of archaea and bacteria. PLoS biology, 21(4), e3002083.

Roux, S., Emerson, J. B., Eloe-Fadrosh, E. A., & Sullivan, M. B. (2017). Benchmarking viromics: an in silico evaluation of metagenome-enabled estimates of viral community composition and diversity. PeerJ, 5, e3817. doi:10.7717/peerj.3817

Roux, S., Enault, F., Hurwitz, B. L., & Sullivan, M. B. (2015). VirSorter: mining viral signal from microbial genomic data. PeerJ, 3, e985. doi:10.7717/peerj.985

Roux S., Solonenko N. E., Dang V. T., Poulos B. T., Schwenck S. M., Goldsmith D. B., Coleman M. L., Breitbart M., Sullivan M. B. (2016). Towards quantitative viromics for both double-stranded and single-stranded DNA viruses. PeerJ, 4, e2777. doi:10.7717/peerj.2777

Roux S., Trubl G., Goudeau D., Nath N., Couradeau E., Ahlgren N. A., Zhan Y., Marsan D., Chen F., Fuhrman J. A., Northen T. R., Sullivan M. B., Rich V. I., Malmstrom R. R. & Eloe-Fadrosh, E. A. (2019a). Optimizing de novo genome assembly from PCR-amplified metagenomes. PeerJ, 7, e6902.

Santos-Medellin, C., Zinke, L. A., Ter Horst, A. M., Gelardi, D. L., Parikh, S. J., & Emerson, J. B. (2021). Viromes outperform total metagenomes in revealing the spatiotemporal patterns of agricultural soil viral communities. The ISME Journal. doi:10.1038/s41396-021-00897-y

Sato, M. P., Ogura, Y., Nakamura, K., Nishida, R., Gotoh, Y., Hayashi, M., Hisatsune, J., Sugai, M., Takehiko, I. & Hayashi, T., (2019). Comparison of the sequencing bias of currently available library preparation kits for Illumina sequencing of bacterial genomes and metagenomes. DNA Research, 26(5), 391–398.

Schmidt, O., Hink, L., Horn, M. A., & Drake, H. L. (2016). Peat: home to novel syntrophic species that feed acetate- and hydrogen-scavenging methanogens. The ISME Journal, 10(8), 1954–1966. doi:10.1038/ismej.2015.256

Schostag, M., Priemé, A., Jacquiod, S., Russel, J., Ekelund, F., & Jacobsen, C. S. (2019). Bacterial and protozoan dynamics upon thawing and freezing of an active layer permafrost soil. The ISME Journal, 13(5), 1345–1359. doi:10.1038/s41396-019-0351-x

Schuur, E. A., & Mack, M. C. (2018). Ecological response to permafrost thaw and consequences for local and global ecosystem services. Annual review of ecology, evolution, and systematics, 49, 279–301.

Shaffer, M., Borton, M. A., McGivern, B. B., Zayed, A. A., La Rosa, S. L., Solden, L. M., Liu, P., Narrowe, A. B., Rodríguez-Ramos, J., Bolduc, B., Gazitúa, M. C., Daly, R. A., Smith, G. J., Vik, D. R., Pope, P. B., Sullivan, M. B., Roux, S. & Wrighton, K. C. (2020). DRAM for distilling microbial metabolism to automate the curation of microbiome function. Nucleic acids research, 48(16), 8883–8900.

Shmakova, L. A., & Rivkina, E. M. (2015). Viable eukaryotes of the phylum Amoebozoa from the Arctic permafrost. Paleontological Journal, 49, 572–577.

Shukla, P. R., Skea, J., Calvo Buendia, E., Masson-Delmotte, V., Pörtner, H. O., Roberts, D. C., Zhai, P., Slade, R., Connors, S., Van Diemen, R., & Ferrat, M. (2019). IPCC, 2019: Climate Change and Land: an IPCC special report on climate change, desertification, land degradation, sustainable land management, food security, and greenhouse gas fluxes in terrestrial ecosystems.

Singleton, C. M., McCalley, C. K., Woodcroft, B. J., Boyd, J. A., Evans, P. N., Hodgkins, S. B., Chanton, J. P., Frolking, S., Crill, P. M., Saleska, S. R., Rich, V. I. & Tyson, G. W. (2018). Methanotrophy across a natural permafrost thaw environment. The ISME journal, 12(10), 2544–2558.

Solonenko, S. A., Ignacio-Espinoza, J. C., Alberti, A., Cruaud, C., Hallam, S., Konstantinidis, K., Tyson, G., Wincker, P. & Sullivan, M. B. (2013a). Sequencing platform and library preparation choices impact viral metagenomes. BMC Genomics, 14(1), 320. doi:10.1186/1471-2164-14-320

Solonenko, S. A., & Sullivan, M. B. (2013b). Preparation of metagenomic libraries from naturally occurring marine viruses. Methods in enzymology, 531, 143–165.

Sommers, P., Fontenele, R. S., Kringen, T., Kraberger, S., Porazinska, D. L., Darcy, J. L., Schmidt, S. K. & Varsani, A. (2019). Single-stranded DNA viruses in antarctic cryoconite holes. Viruses, 11(11), 1022.

Starr, E. P., Shi, S., Blazewicz, S. J., Koch, B. J., Probst, A. J., Hungate, B. A., Pett-Ridge, J., Firestone, M. K. & Banfield, J. F. (2021). Stable-isotope-informed, genome-resolved metagenomics uncovers potential cross-kingdom interactions in rhizosphere soil. Msphere, 6(5), e00085–21.

Suttle, C. A. (2005). Viruses in the sea. Nature, 437(7057), 356–361.

Suttle, C. A. (2007). Marine viruses—major players in the global ecosystem. Nature reviews microbiology, 5(10), 801–812.

Tan, B., Charchuk, R., Li, C., Nesbø, C., Abu Laban, N., & Foght, J. (2014). Draft genome sequence of uncultivated Firmicutes (Peptococcaceae SCADC) single cells sorted from methanogenic alkane-degrading cultures. Genome announcements, 2(5), e00909–14.

Ter Horst, A. M., Santos-Medellín, C., Sorensen, J. W., Zinke, L. A., Wilson, R. M., Johnston, E. R., Trubl, G., Pett-Ridge, J., Blazewicz, S. J., Hanson, P. J., Chanton, J. P., Schadt, C. W., Kostka, J. E. & Emerson, J. B. (2020). Minnesota peat viromes reveal terrestrial and aquatic niche partitioning for local and global viral populations. Cold Spring Harbor Laboratory.

Thompson, L. R., Sanders, J. G., McDonald, D., Amir, A., Ladau, J., Locey, K. J., Prill, R. J., Tripathi, A., Gibbons, S. M., Ackermann, G., Navas-Molina, J.A., Janssen, S., Kopylova, E., Vázquez-Baeza, Y., González, A., Morton, J. T., Mirarab, S., Xu, Z. Z., Jiang, L., Haroon, M. F., Kanbar, J., Zhu, Q., Song, S. J., Kosciolek, T., Bokulich, N. A., Lefler, J., Brislawn, C. J., Humphrey, G., Owens, S. M., Hampton-Marcell, J., Berg-Lyons, D., McKenzie, V., Fierer, N., Fuhrman, J. A., Clauset, A., Stevens, R. L., Shade, A., Pollard, K. S., Goodwin, K. D., Jansson, J. K., Gilbert, J. A., Knight, R. & The Earth Microbiome Project Consortium (2017). A communal catalogue reveals Earth’s multiscale microbial diversity. Nature, 551(7681), 457–463.

Trubl, G., Hyman, P., Roux, S., & Abedon, S. T. (2020). Coming-of-Age Characterization of Soil Viruses: A User’s Guide to Virus Isolation, Detection within Metagenomes, and Viromics. Soil Systems, 4(2), 23. doi:10.3390/soilsystems4020023

Trubl, G., Jang, H. B., Roux, S., Emerson, J. B., Solonenko, N., Vik, D. R., Solden, L., Ellenbogen, J., Runyon, A. T., Bolduc, B., Woodcroft, B.J., Saleska, S. R., Tyson, G. W., Wrighton, K. C., Sullivan, M. B. & Rich, V. I. (2018). Soil Viruses Are Underexplored Players in Ecosystem Carbon Processing. mSystems, 3(5). doi:10.1128/msystems.00076-18

Trubl, G., Kimbrel, J. A., Liquet-Gonzalez, J., Nuccio, E. E., Weber, P. K., Pett-Ridge, J., Jansson, J. K., Waldrop, M. P. & Blazewicz, S. J. (2021). Active virus-host interactions at sub-freezing temperatures in Arctic peat soil. Microbiome, 9(1), 1–15.

Trubl, G., Roux, S., Solonenko, N., Li, Y. F., Bolduc, B., Rodríguez-Ramos, J., Eloe-Fadrosh, E. A., Rich, V. I. & Sullivan, M. B. (2019). Towards optimized viral metagenomes for double-stranded and single-stranded DNA viruses from challenging soils. PeerJ, 7, e7265.

Van Goethem, M. W., Swenson, T. L., Trubl, G., Roux, S., & Northen, T. R. (2019). Characteristics of Wetting-Induced Bacteriophage Blooms in Biological Soil Crust. mBio, 10(6). doi:10.1128/mbio.02287-19

Ward, C. P., Nalven, S. G., Crump, B. C., Kling, G. W., & Cory, R. M. (2017). Photochemical alteration of organic carbon draining permafrost soils shifts microbial metabolic pathways and stimulates respiration. Nature communications, 8(1), 772.

Wawrik, B., Marks, C. R., Davidova, I. A., McInerney, M. J., Pruitt, S., Duncan, K. E., Suflita, J. M. & Callaghan, A. V. (2016). Methanogenic paraffin degradation proceeds via alkane addition to fumarate by ‘Smithella’spp. mediated by a syntrophic coupling with hydrogenotrophic methanogens. Environmental Microbiology, 18(8), 2604–2619. doi:10.1111/1462-2920.13374

Williamson, K. E., Fuhrmann, J. J., Wommack, K. E., & Radosevich, M. (2017). Viruses in soil ecosystems: an unknown quantity within an unexplored territory. Annual review of virology, 4, 201–219. X

Wilson, R. M., Hough, M. A., Verbeke, B. A., Hodgkins, S. B., Tyson, G., Sullivan, M. B., Brodie, E., Riley, W. J., Woodcroft, B. J., McCalley, C., Dominguez, S. C., Crill, P. M., Varner, R. K., Frolking, S., Cooper, W. T., Chanton, J. P., Saleska, S. R., Rich, V. I. & Tfaily, M. M. (2022). Plant organic matter inputs exert a strong control on soil organic matter decomposition in a thawing permafrost peatland. Science of The Total Environment, 820, 152757.

Woodcroft, B. J., Singleton, C. M., Boyd, J. A., Evans, P. N., Emerson, J. B., Zayed, A. A., Hoelzle, R. D., Lamberton, T. O., McCalley, C. K., Hodgkins, S. B., Wilson, R.M., Purvine, S. O., Nicora, C. D., Li, C., Frolking, S., Chanton, J. P., Crill, P. M., Saleska, S. R., Rich, V. I. & Tyson, G. W. (2018). Genome-centric view of carbon processing in thawing permafrost. Nature, 560(7716), 49–54. doi:10.1038/s41586-018-0338-1

Wu, M. H., Chen, S. Y., Chen, J. W., Xue, K., Chen, S. L., Wang, X. M., Chen, T., Kang, S. C., Rui, J. P., Thies, J. E., Bardgett, R. D. & Wang, Y. F. (2021). Reduced microbial stability in the active layer is associated with carbon loss under alpine permafrost degradation. Proceedings of the National Academy of Sciences, 118(25), e2025321118.

Wu, R., Smith, C. A., Buchko, G. W., Blaby, I. K., Paez-Espino, D., Kyrpides, N. C., Yoshikuni, Y., McDermott, J. E., Hofmockel, K. S., Cort, J. R. & Jansson, J.K. (2022). Structural characterization of a soil viral auxiliary metabolic gene product–a functional chitosanase. Nature Communications, 13(1), 1–14.

Zawar-Reza, P., Argüello-Astorga, G. R., Kraberger, S., Julian, L., Stainton, D., Broady, P. A., & Varsani, A. (2014). Diverse small circular single-stranded DNA viruses identified in a freshwater pond on the McMurdo Ice Shelf (Antarctica). Infection, Genetics and Evolution, 26, 132–138.

Zhao, L., Lavington, E., & Duffy, S. (2021). Truly ubiquitous CRESS DNA viruses scattered across the eukaryotic tree of life. Journal of Evolutionary Biology, 34(12), 1901–1916. doi:10.1111/jeb.13927

Zimmerman, A. E., Howard-Varona, C., Needham, D. M., John, S. G., Worden, A. Z., Sullivan, M. B., Waldbauer, J. R. & Coleman, M. L. (2020). Metabolic and biogeochemical consequences of viral infection in aquatic ecosystems. Nature Reviews Microbiology, 18(1), 21–34. doi:10.1038/s41579-019-0270-x

