## Supplementary figures and images for "Population ecology and potential biogeochemical impacts of ssDNA and dsDNA soil viruses along a permafrost thaw gradient"

### Supplementary Figure 1

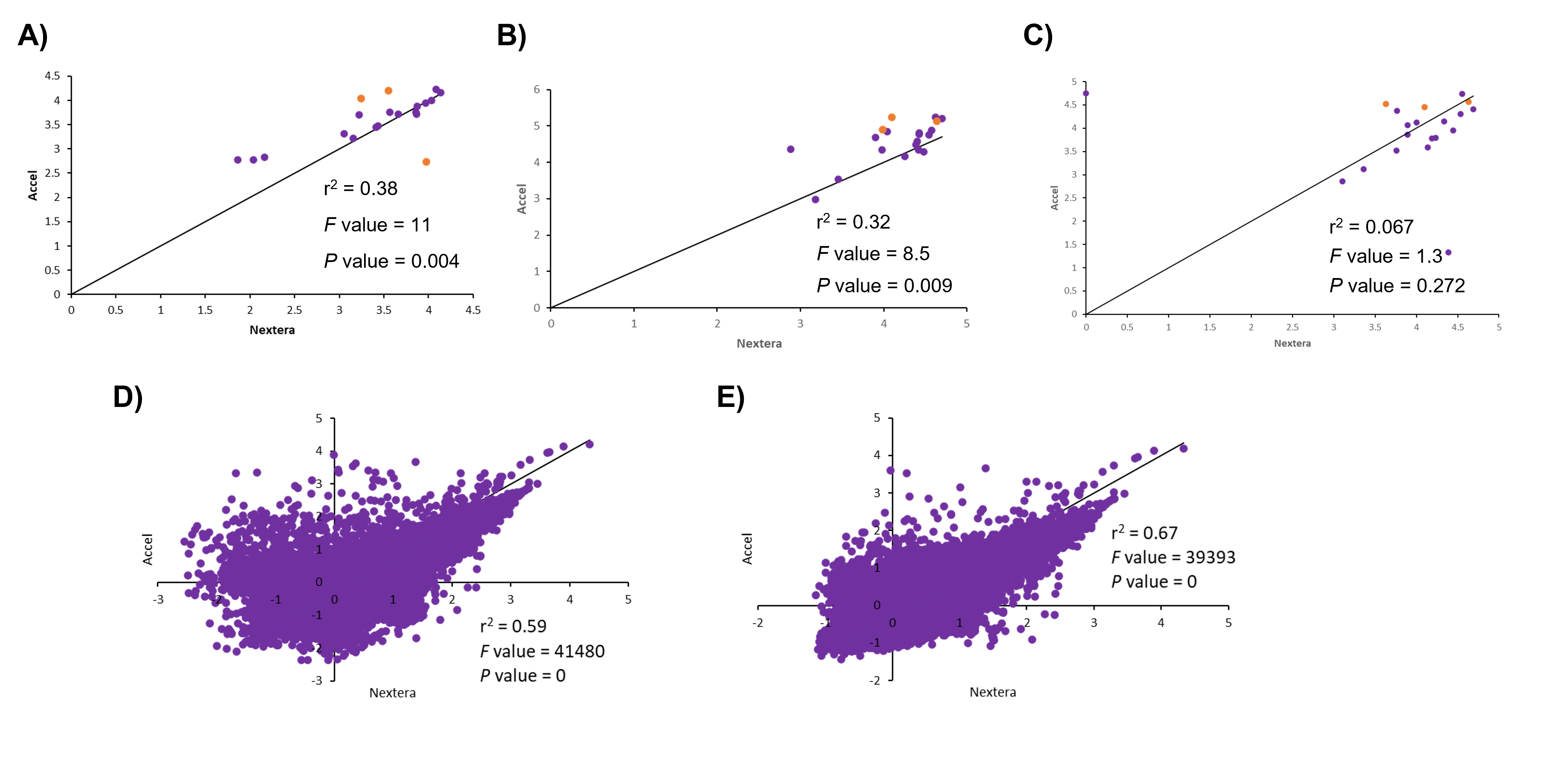

### Supplementary Figure 2

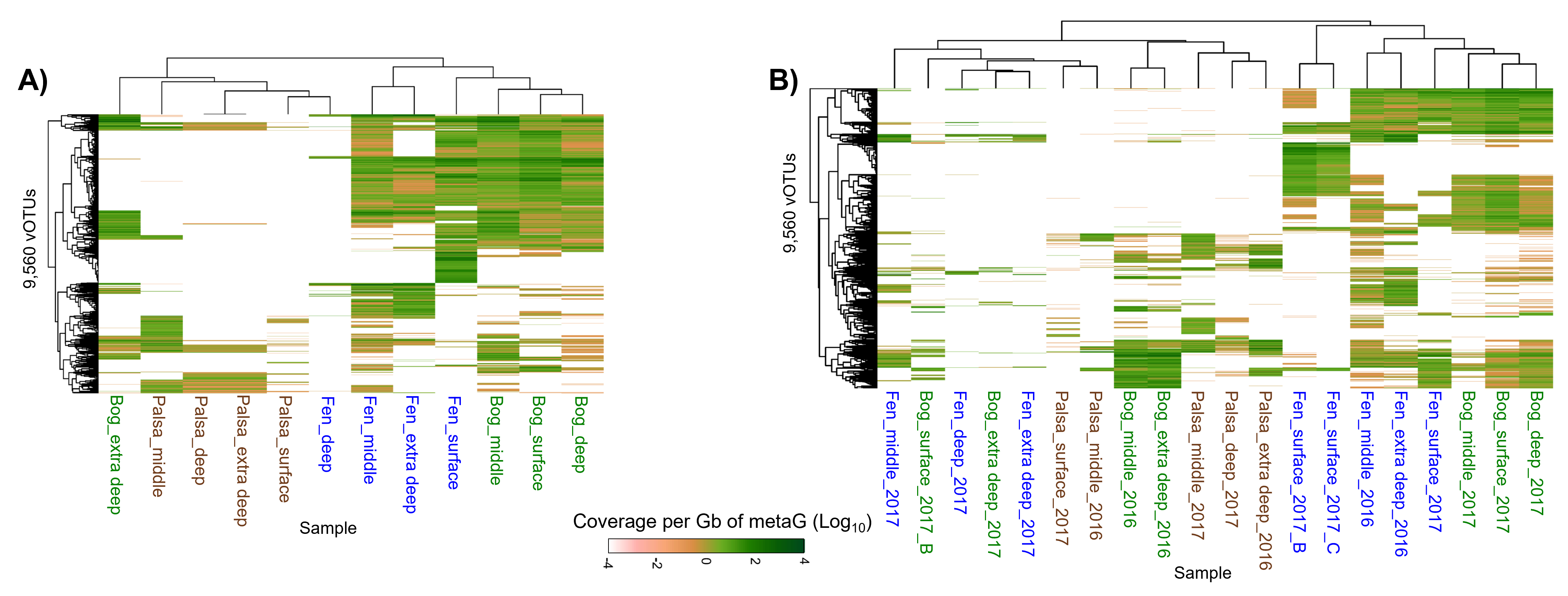
